# Mechanistic insights and clinical implications of cross-reactive anti-prophage antibodies and bacterial heteroresistance on phage therapeutic failure

**DOI:** 10.1101/2025.04.22.650118

**Authors:** F Gordillo Altamirano, D Subedi, M Beiers, M Bucher, S Dahlman, DM Patel, M Parker, D Korneev, MJ Robinson, K Pragastis, J Wisniewski, C Rees, H Ramshaw, SF Khan, BJ Gardiner, Y Hammerschlag, D Keating, T Kotsimbos, J Hawkey, JJ Barr, AY Peleg

## Abstract

Phage therapy is an exciting strategy against antimicrobial-resistant bacterial infections, but critical knowledge gaps regarding its clinical application persist. Studying a patient with a life-threatening, chronic bacterial infection who failed phage therapy, we uncovered important biological concepts with direct translational impact. Using longitudinal clinical samples, we found that patients can harbour pre-existing antibodies against active prophages induced from the genome of the causative pathogen. Notably, these antibodies can contribute to clinical failure by cross-reacting with and effectively neutralising therapeutic phage. We also uncovered bacterial heteroresistance, characterised by bacterial subpopulations from the initial infection with reduced phage susceptibility, as a further contributor to treatment failure. These findings highlight the intricate interplay between host immunology, bacterial genetic diversity and phage biology, bearing broad significance for clinical phage therapy. Future phage therapy patients, especially those with chronic infections, should be screened for antiphage immunity and bacterial heteroresistance prior to phage treatment.

## Introduction

Antimicrobial resistance (AMR) is one of the greatest health threats we currently face. The latest global estimations suggest that in 2021, bacterial AMR directly caused over a million deaths, while playing a role in just under 5 million ^1^. AMR is not only a leading cause of death but poses a significant burden on the global economy and healthcare systems ^2^. These consequences are expected to increase unless a concerted, multi-disciplinary effort is deployed to fight AMR, including the development of therapeutic strategies for when antibiotics prove ineffective, such as phage therapy.

Phage therapy is the clinical use of bacteriophages, viruses that specifically kill bacteria, to treat bacterial infections ^3^. Although phages were discovered and used before antibiotics, phage therapy is now experiencing a resurgence amidst the AMR crisis. In the last two decades, we have accumulated a significant body of evidence from preclinical studies suggesting that phage therapy is safe and potentially effective ^4^, encouraging translational efforts. As for clinical use, a recent systematic review examined 2,241 patients from 59 studies and deemed phage therapy a promising strategy in the fight against difficult-to-treat infections and AMR ^5^. However, most of this clinical experience comes from case reports and case-series, with only a handful of clinical trials. Substantial knowledge gaps remain to be addressed before phage therapy becomes standardised and widespread, including questions around the pharmacology of phages, their interactions with antibiotics, and the tripartite relationship between phages, bacteria and the human host ^6^.

We present a patient with a chronic, extensively drug-resistant (XDR) bacterial (*Bordetella bronchialis*) infection associated with life-threatening infective pulmonary exacerbations and bacteraemia that was poorly controlled with traditional antibiotics. The patient qualified for compassionate use phage therapy. We report the isolation and detailed characterisation of the therapeutic phage, and the clinical course of the patient that resulted in phage therapeutic failure with bacteriological relapse. We explored the complex *in vivo* dynamics of host immunology, bacteriology, and phage biology, and have provided essential clinical insights into the future treatment of life-threatening bacterial infections with bacteriophages.

## Results and Discussion

### Recurrent life-threatening infection with extensively drug-resistant bacteria

A 22-year-old male patient with cystic fibrosis (CF) was referred to the Victorian Bacteriophage Therapy Program (VIC*Phage*) in 2022 for compassionate use phage treatment of recurrent, invasive *Bordetella bronchialis* infection, including pneumonia and bacteraemia. He had a history of a liver-pancreas transplant six years prior for CF-related liver disease and diabetes mellitus, and end-stage kidney disease secondary to calcineurin inhibitor use, aminoglycoside exposure, and diabetes mellitus. He was on maintenance immunosuppression with tacrolimus and prednisone.

*B. bronchialis* was first isolated from the patient’s sputum at the age of 10 years. *B. bronchialis*, known before 2015 as *Bordetella* genogroup 3, are clinically uncommon motile Gram-negative bacilli that are facultatively aerobic and slow-growing ^7^. Despite reported cases of this pathogen in patients with CF ^8,9^, there is limited information regarding its pathogenesis, biology, or epidemiology. Through his teenage years, the patient developed recurrent infective respiratory exacerbations with repeated hospitalisation and isolation of *B. bronchialis*. Initially, the isolate tested susceptible to ceftazidime and piperacillin-tazobactam (Table S1). However, repeated infections ensued and in the three years prior to referral, he was admitted nine times for infective exacerbations, including three episodes of invasive bacteraemia, an uncommon complication in patients with CF. To prevent or reduce the frequency of these life-threatening exacerbations, ongoing minocycline and trimethoprim-sulfamethoxazole were commenced in 2021.

In June 2022, the patient presented with fever, productive cough, and dyspnoea. Breakthrough *B. bronchialis* was isolated from sputum and multiple sets of blood cultures. Chest imaging demonstrated diffuse, severe bilateral bronchiectasis with extensive mucous plugging and patchy foci of consolidation. He was commenced on empiric IV meropenem and inhaled colistin, and antimicrobial susceptibility testing demonstrated that the isolate had become XDR (Table S1, Note S1). At this point, referral for compassionate use phage treatment was received.

### Isolation and characterisation of an active phage

Using wastewater samples from Melbourne, Australia, and cultures of a single *B. bronchialis* colony obtained from the patient’s sputum in June 2022 (wild type [wt] *B. bronchialis*), we successfully discovered, isolated and amplified phage øSimón (Fig. 1a). Multiple efforts to isolate additional phages were unsuccessful. In liquid coculture, øSimón inhibited bacterial growth for up to 40 hours at multiplicities of infection (MOI) as low as 0.01 (Fig. 1b), and consistently achieved titres above 10^8^ plaque forming units (pfu)/ml. In a one-step kill curve, we determined that øSimón had a latent period of ~2.5 h, which is more than three times faster than the calculated doubling time for the host strain (8.3 ± 2.6 h), and a burst size of nearly 30 virions per cell (Fig. 1c). øSimón displayed similar activity against five other clinical *B. bronchialis* isolates (three from blood, two from sputum) obtained from the patient prior to starting phage therapy (Fig. 1d). To mimic the more hostile environment of CF sputum, we also assessed øSimón’s activity in artificial sputum media (ASM). The bactericidal activity of øSimón in ASM was slower, less profound, but more sustained than in liquid culture (Fig. 1e), while also resulting in slightly higher phage titres (Fig. 1f). Transmission electron microscopy (TEM) showed that øSimón was morphologically a siphovirus, with a total length of ~226 nm (Fig. 1g). øSimón’s genome did not encode any markers of lysogeny, AMR, toxins or virulence (Fig. 1h, Note S2), making it an ideal candidate for therapeutic use.

**Figure 1.**
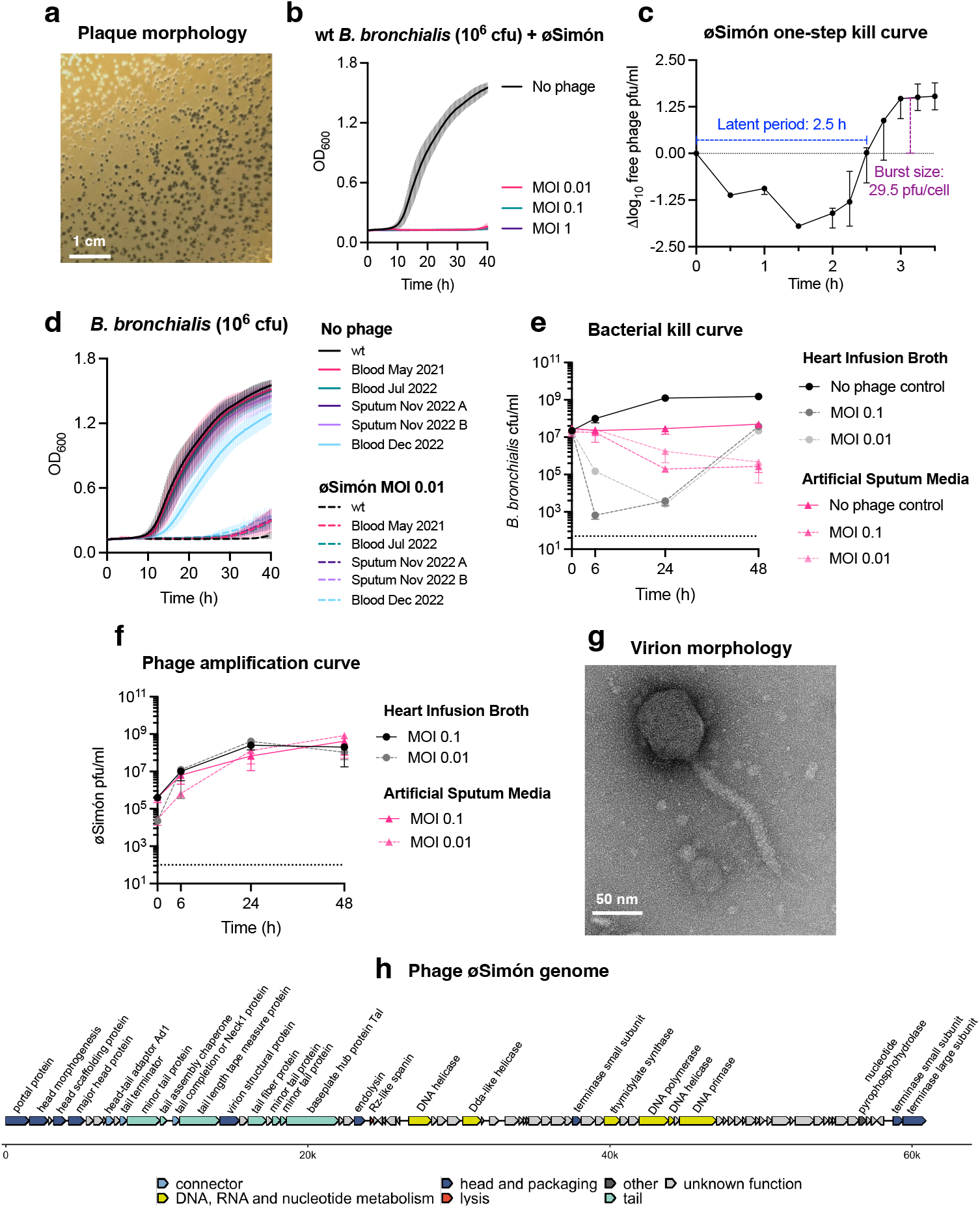
Characterisation of phage øSimón. **a:** Plaque morphology on HI agar, 48 h of incubation. Scale bar: 1 cm. **b:** Optical density at 600 nm (OD_600_) growth curve (mean ± SD; n = 3) of *B. bronchialis* without phage (black) and with three different multiplicities of infection (MOI) of øSimón. **c:** One-step kill curve (median ± 95% CI; n = 3). **d:** Sensitivity of further *B. bronchialis* isolates from the patient against øSimón in liquid culture (growth curve, mean ± SD; n = 3). Isolates are labelled by type of clinical sample and date of isolation. **e and f**: Comparison of øSimón’s bactericidal activity and amplification in liquid (heart infusion broth, black) and artificial sputum media (magenta) (median ± 95% CI; n = 3; dotted lines represent the limit of detection). **g:** øSimón virion under transmission electron microscopy. Scale bar: 50 nm. **h:** Assembled øSimón genome (60,921 bp, 80 open reading frames).

### Clinical administration of øSimón

A clinical-grade preparation of øSimón was produced locally at the Monash Phage Foundry ^10^. All product batches (Table S2) underwent internal and third-party validation for phage titre, sterility, and endotoxin levels suitable for IV administration ^11^. In the setting of a repeated episode of invasive bacteraemia with XDR *B. bronchialis*, the patient received the first dose of IV øSimón (5.4 × 10^7^ pfu/dose) (day 0) (Fig. 2). Phage therapy was administered twice daily and, given the severity of infection, was co-administered alongside IV meropenem and inhaled colistin. The patient stabilised clinically, with blood cultures clearing by day 5. After close clinical and laboratory monitoring for six days, the patient was transferred to our hospital-in-the-home program for ongoing IV phage management, completing 30 days of treatment ^12^. Phage therapy was well tolerated, associated with a reduction in inflammatory markers (C-reactive protein) (Fig. S1), and no clinical or laboratory adverse effects were reported.

**Figure 2.**
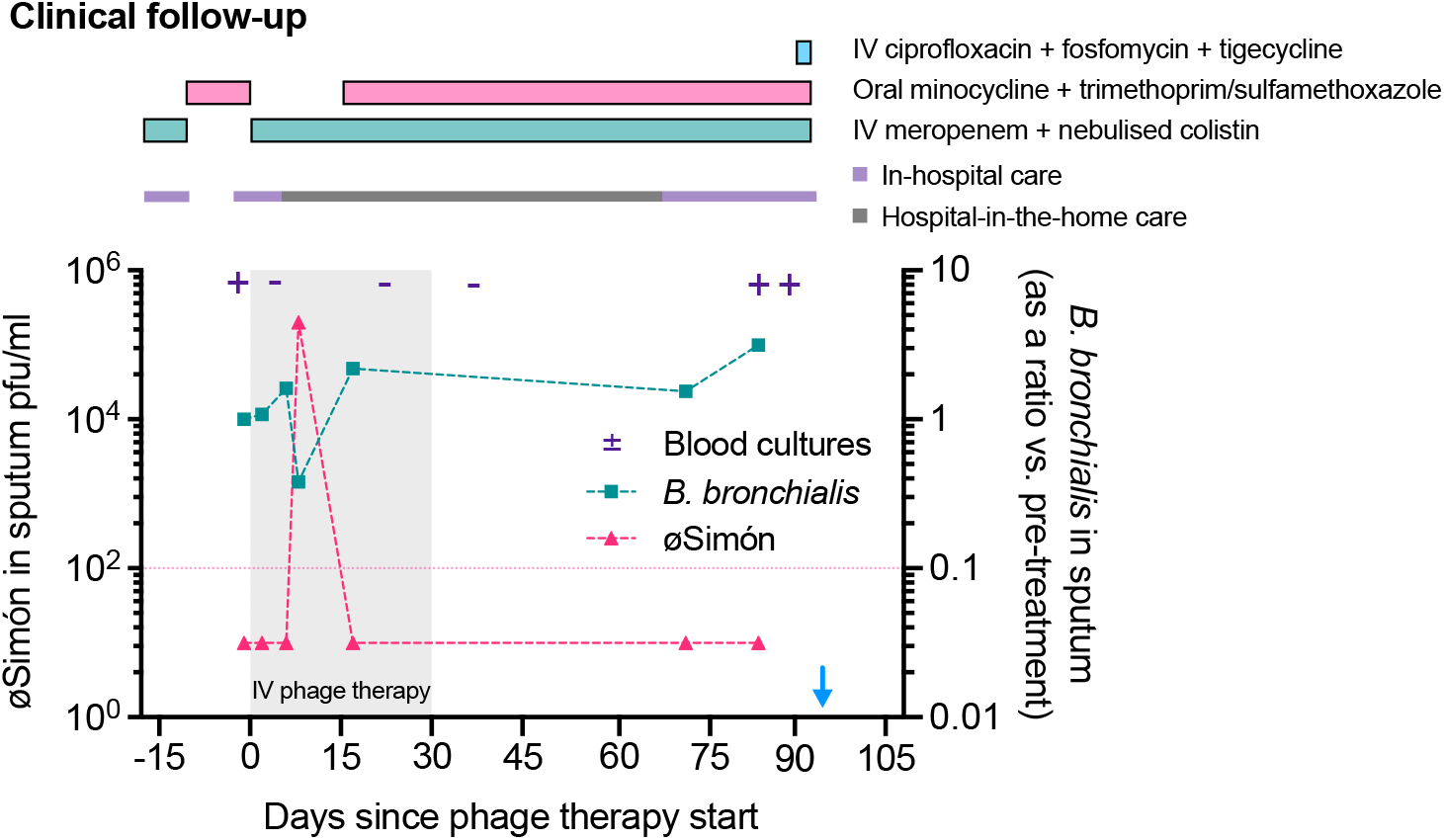
Clinical follow-up of phage therapy. The patient received the first dose of phage therapy on Day 0 (Dec 20^th^, 2022) and passed away on day 93 (blue arrow). The grey shade in the plot represents the 30-day period of IV phage therapy. Laboratory tests included blood cultures for *B. bronchialis* (positive [+] and negative [-] purple symbols), and quantification of *B. bronchialis* (right y-axis) and øSimón (left y-axis) in the patient’s sputum. Bacterial load is presented as a ratio of the pre-treatment amount (1.3 × 10^8^ cfu/ml). Dotted line highlights the limit of detection for øSimón. Atop the graph, coloured rectangles denote antibiotic therapy administered to the patient, and the coloured line differentiates between inpatient and outpatient care.

Alongside these clinical outcomes, we observed an almost 80% reduction in the amount of *B. bronchialis* in the patient’s sputum at day 8 of phage treatment, and notably, this coincided with the finding of actively replicating øSimón (Fig. 2). However, by day 17, the *B. bronchialis* load increased back to pre-treatment levels, and øSimón was no longer isolated from the sputum. The patient went on to develop progressive complications of his end-stage renal disease. In the context of a repeat admission for worsening illness (day 68), he developed further episodes of *B. bronchialis* bacteraemia and passed away on day 93, two months after the last dose of øSimón. Given the undulating clinical and microbiological course, we investigated the possible mechanisms of phage therapy failure and the dynamic interactions between bacteria, phage, and human host over time.

### Pre-existing antiphage immunity

Despite phages being non-pathogenic to humans, it is known that the human immune system produces antibodies against them ^13^. Antiphage antibodies can neutralise phage activity ^14^, either directly, by targeting surface-exposed structural proteins involved in bacterial adsorption ^15,16^, or, to a lesser degree, indirectly by opsonising them for phagocytosis ^13,16,17^. However, how much this immune response, or lack thereof, contributes to the outcomes of phage therapy is still debated ^18–21^. The humoral response is T-cell dependent, and activated B-cells differentiate into plasma cells to produce a specific, polyclonal, antiphage antibody response ^13,22^. Circulating antibodies usually peak one to two weeks after initial contact with a phage ^13,23^, aligning with the time we observed a reduction of phage activity in our patient.

Using patient serum samples from days 0, 7, and 21, we first performed a phage neutralisation assay ^21^. Briefly, we tested the ability of three successive 10-fold serum dilutions to neutralise a known dose of øSimón in 30 min. We used an unrelated, anti-*Pseudomonas aeruginosa* phage (øFaith) as a control under the same conditions. Notably, at day 0, before the patient had received the first dose of øSimón, the serum could neutralise ~25% of the phage dose (Fig. 3a). The neutralisation increased to ~60% on days 7 and 21. No neutralisation was seen against the control phage. We also used a more standardised metric known as the phage inactivation rate (*k*), which generates “high”, “moderate”, and “low” thresholds, inversely correlating with odds of treatment success ^19,21^. We showed that the patient’s pre-treatment serum had a “moderate” *k* value of 10.69, which increased to “high” at days 7 and 21 (27.67 and 30.60, respectively) (Fig. 3b). Notably, we also observed low phage neutralisation capacity in the patient’s sputum (Fig. 3c and 3d).

**Figure 3.**
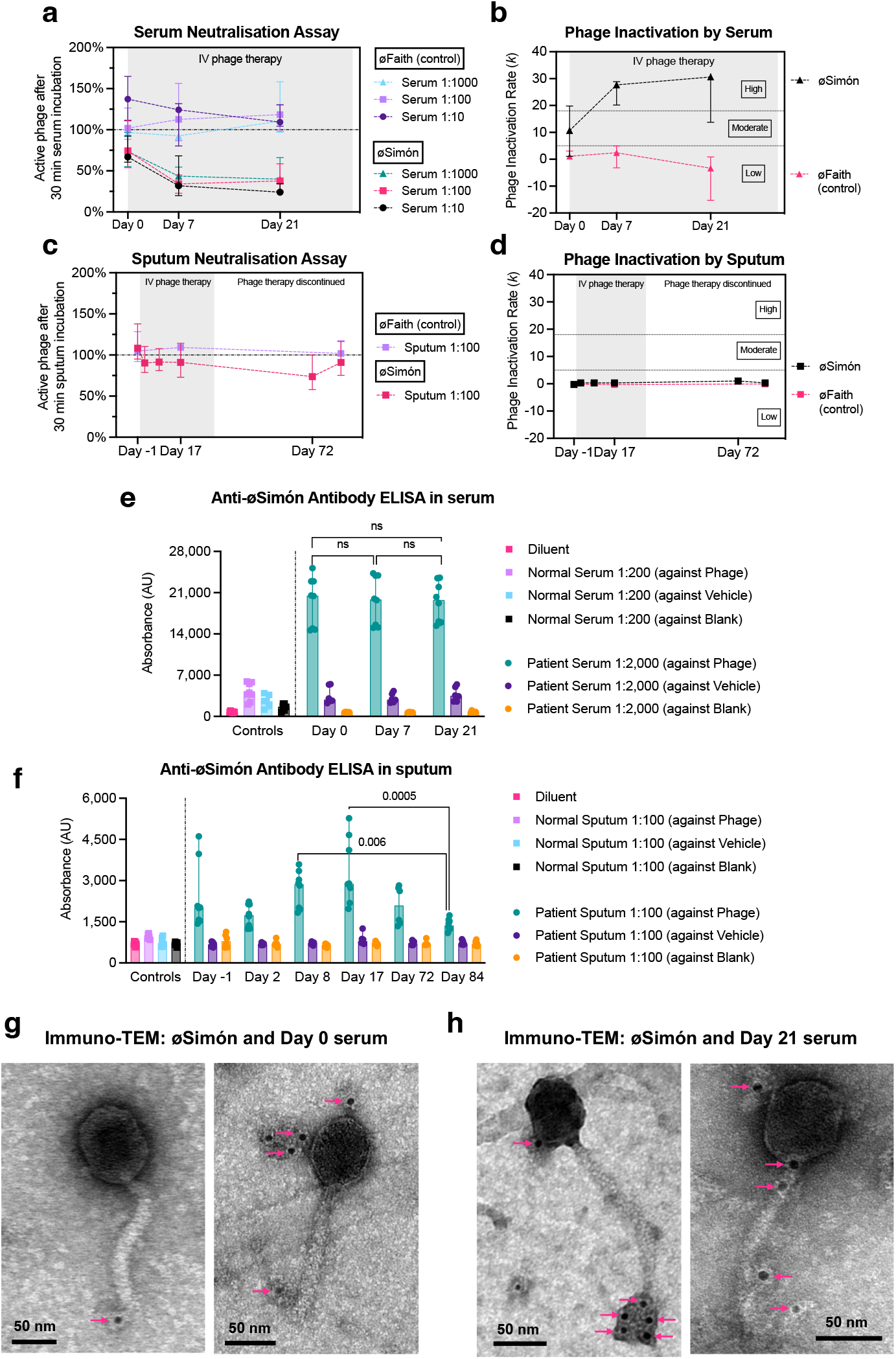
The patient harboured pre-existing antiphage immunity. **a:** Serum neutralisation assay (10^6^ pfu). The x-axis follows the treatment timeline, and the y-axis plots the percentage of active phage (soft overlay agar assay) (median ± 95% CI; n = 3). **b:** Calculation of phage inactivation rate (*k*) with data from the previous assay (median ± 95% CI; n = 3). **c and d:** The experiments depicted in a and b were repeated using patient’s filter-sterilised sputum instead of serum. **e and f:** Indirect ELISA looking for anti-øSimón antibodies in serum (e) and sputum (f). Controls were run with serum from a healthy donor or sputum from a patient with cystic fibrosis (median ± 95% CI; all technical replicates from two experiments plotted; Kruskal-Wallis test; if not plotted the comparison was not statistically significant). **g and h:** Immuno-TEM micrographs of øSimón with pre- and post-treatment patient serum, respectively, and gold-labelled anti-human IgG detection antibodies (black dots with magenta arrowheads). Scale bars: 50 nm. Additional micrographs and controls in Fig. S2.

To confirm that the phage inactivation was mediated by anti-øSimón antibodies, we designed an indirect ELISA assay. Intact, highly purified ^24^ øSimón was used as the coating antigen. To further discriminate the presence of confounding antibodies against residual endotoxin or other bacterial products, we also used a vehicle control obtained by filtering out (0.02 μm) phage particles from the coating product. Sera from the patient were run in parallel with a serum control from a healthy patient, and all tests used an isotype pan-specific detection antibody. The ELISA confirmed the presence of anti-øSimón antibodies at days 0, 7, and 21 (Fig. 3e). Anti-øSimón antibodies were also detected in the patient’s sputum, although the signal was markedly less intense than in serum (Fig. 3f). Interestingly, a significant peak of antibody was detected in the sputum on days 8 and 17, while the patient was receiving IV phage therapy, compared to follow-up sputum from day 84, almost two months after phage therapy had stopped.

We next sought to visualise the phage structures being targeted by the antibodies. To do this, we purified the IgG fraction from the patient’s serum and used it in an immuno-TEM experiment with øSimón and gold-labelled anti-human IgG detection antibodies. We observed that the patient’s antibodies were binding to multiple sites on the virion, including the capsid, neck, tail, and tail fibres (Fig. 3g, 3h and S2). To further confirm the role of these anti-øSimón antibodies in phage inactivation, we serially passaged the day 0 serum sample on ELISA wells coated with øSimón to deplete the serum of anti-øSimón antibodies (Fig. S3a, Note S3). We repeated the neutralisation assay with this depleted serum and observed that the neutralisation effect was reversed (Fig. S3b). We also ruled out that the neutralisation effect was mediated by antibodies targeting the bacterial surface and inhibiting phage adsorption, as pre-treatment of the bacteria with the patient’s serum had no effect on øSimón adsorption (Fig. S4a and S4b). To summarise, we demonstrated the presence of antibodies capable of recognising and inactivating øSimón in both serum and sputum of the patient even prior to phage therapy. These antibody levels were maintained throughout the duration of therapy and beyond, but their neutralising effect increased over time.

### Resident prophages as drivers of pre-existing immunity to therapeutic phage

We investigated a possible explanation for the presence of pre-existing anti-øSimón antibodies. Given the seemingly infinite diversity of phages in the biosphere ^25^, we believe it was unlikely for the patient to have previously encountered øSimón. However, as the patient had harboured the *B. bronchialis* infection for over a decade, we hypothesised he may have been exposed to other phages with activity against the pathogen, sharing similar structural features to øSimón, and triggering a cross-reactive immune response. Furthermore, recent studies using cell-free DNA sequencing have demonstrated that patients with sepsis have an overrepresentation of pathogen-specific phages ^26,27^. This could be explained by prophages, which are phages that have integrated into a bacterium’s chromosome and occasionally re-enter the lytic cycle in periods of rapid bacterial expansion or stress ^27^. Consequently, we sought to investigate whether prophages integrated into the patient’s *B. bronchialis*, and reactivated at different times during the chronic infection, could have been responsible for cross-reactive immunity against øSimón.

We parsed the genome of the wt *B. bronchialis* for prophage regions. We identified three prophages with medium-to-high completeness scores ^28^. On a nucleotide level, they did not show homology to øSimón (Fig. 4a, S5a). Next, we explored whether these prophages could re-enter the lytic lifecycle by the addition of an induction agent (mitomycin C) to a bacterial liquid culture, or spontaneously in the absence of an induction agent. After purification ^24^, we confirmed the presence of phage virions through TEM and DNA sequencing, but only in the culture with mitomycin C. TEM micrographs of the suspension detected phage particles at a concentration of ~10^6^ virions/ml, with a single (siphovirus) morphology (Fig. 4b). After sequencing, two thirds of the obtained reads mapped to prophage region 2, a 51.5 kb-long sequence with the closest match being *Burkholderia* phage Bups phi1 (GenBank accession numbers EU307292.1-EU307295.1), which was also a siphovirus. This purified lysate, containing at least one of the induced prophages from the *B. bronchialis* host, was then used for subsequent experiments.

**Figure 4.**
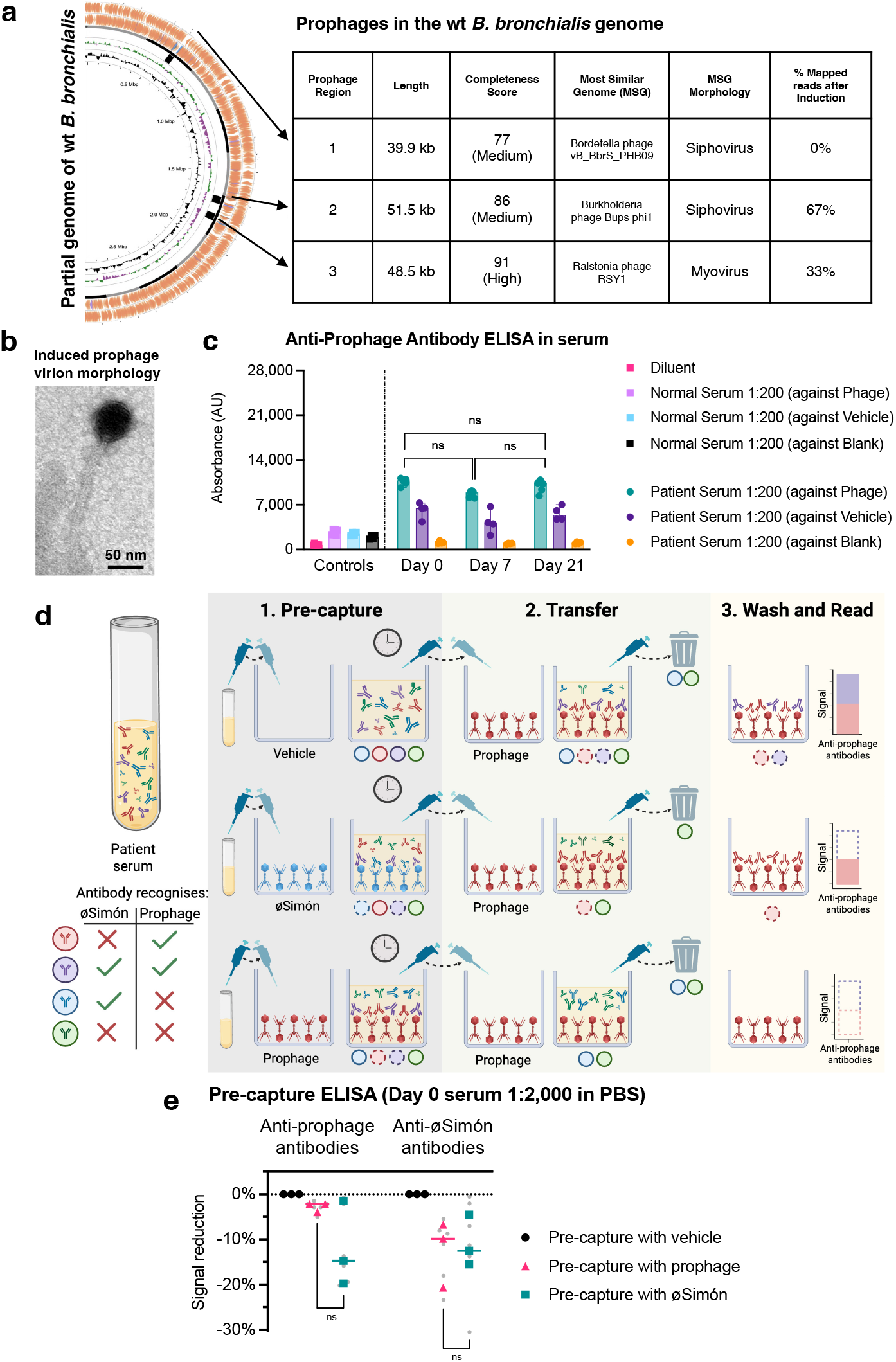
Cross-reactive antibodies between *Bordetella bronchialis* prophage and øSimón. **a:** Screening of prophage regions in the genome of wt *B. bronchialis*. Rings from innermost to outermost: genome length, GC%, GC skew and open reading frames. The adjacent chart provides information about the three prophages found. **b:** Transmission electron micrograph of the virions found after a culture of *B. bronchialis* was treated with mitomycin C for prophage induction. Scale bar = 50 nm. **c:** Indirect ELISA looking for anti-prophage antibodies in serum; (median ± 95% CI; all technical replicates from two experiments plotted; Kruskal-Wallis test; ns: not significant). **d:** Experimental setup for the pre-capture ELISA. Hypothetical antibodies colour-coded by their ability to recognise øSimón (blue), prophage (red), both (purple), or neither (green). Pre-capture step (1), serum is passed through wells that have been coated with either vehicle (blank, top), øSimón (blue virions, middle), or prophage (red virions, bottom). The circles beneath wells represent the colour-coded antibodies either binding to phages (dashed border) or remaining in suspension (solid border). Next, the serum is transferred to fresh, prophage-coated wells (2), left for incubation and binding, and discarded. All unbound antibodies are washed away, and the signal is read (3). The assay is interpreted by measuring the signal reduction between the top condition and each of the remaining two. Not shown in the figure, the assay was later repeated using øSimón to coat wells in step (2). **e:** Pre-capture ELISA. Percentage reduction was calculated using the reading for the vehicle pre-capture condition as baseline (bars are medians, all replicates from three experiments plotted, with technical replicates as grey circles; Wilcoxon test for comparisons, two-tailed; ns: not significant).

Using the induced prophage suspension, an ELISA was developed to detect anti-prophage antibodies, which were confirmed in the patient’s serum but not sputum (Fig. 4c and S6). Although the antibody signal was weaker than in the assays for anti-øSimón antibodies, the two assays were not directly comparable, as the concentration of virions in the solution used to coat the ELISA wells was 100-times lower for the induced prophage.

We then tested whether the anti-prophage and anti-øSimón antibodies were cross-reactive. To do this, we modified our indirect ELISA by adding a pre-capture step; patient serum was first incubated on wells coated with one of three conditions in parallel: 1) phage vehicle (control), 2) øSimón virions, or 3) induced prophage virions. Following this pre-adsorption, we performed two standard indirect ELISAs: one to detect anti-prophage antibodies (Fig. 4d) and another for anti-øSimón antibodies. Theoretically, pre-adsorption with vehicle should yield the highest signal (baseline), while pre-adsorption with the same phage as the ELISA target should result in the lowest signal due to specific antibody depletion. If pre-adsorption with the opposite phage also reduced signal, this would indicate the presence of cross-reactive antibodies. As shown in Fig. 4e, pre-adsorption with either øSimón or prophage virions led to comparable reductions in antibody signal against both targets (mean reduction for anti-prophage antibodies: 2.8% ± 1% with prophage vs. 12% ± 9.5% with øSimón [p = 0.5]; mean reduction for anti-øSimón antibodies: 10.9% ± 5.7% with øSimón vs. 12.4% ± 7.3% with prophage [p > 0.9]). These results support the presence of serum cross-reactive antibodies, capable of simultaneously recognising øSimón and prophage virions.

Together, our data provides experimental evidence that the human body can produce antibodies against induced prophages from a resident bacterial pathogen. These same antibodies were capable of cross-reacting with the therapeutic øSimón and provide at least a partial explanation for the neutralisation activity of the patient’s pre-treatment serum and the potential impact on øSimón efficacy. An alternative hypothesis that we cannot fully discard is that a different, unknown, immunogen to which the patient had been exposed drove the production of antibodies that simultaneously recognised the prophage and øSimón. Our results complement two important earlier findings. First, Halperin *et al*. showed that patients with acute glomerulonephritis produced antibodies against a secreted, prophage-encoded, streptococcal hyaluronidase ^29^. Second, Fluckiger *et al*. demonstrated T-cell-mediated immunity aimed at a specific tail length tape measure protein from an enterococcal prophage in mice and humans, which cross-reacted with tumour-associated antigens ^30^. Taken together, there is a growing body of evidence showing the relevance of prophages and their interactions with the immune system in various states of human health and disease.

Future endeavours in this area include establishing the precise proteins or epitopes responsible for the observed cross-reactivity. We note, for instance, that at the amino acid level, prophage region 2 has a gene coding for a minor tail protein, with a modest homology to øSimón (29% amino acid identity, 87% query coverage) (Fig. S5). Berkson *et al*. have demonstrated that cross-reactive immunity between two therapeutic myoviruses in a phage cocktail was possibly mediated by tail fibre, tail sheath, and major capsid proteins as antigens ^31^. This would be consistent with our immuno-TEM images (Fig. 3g and 3h) showing antibody binding to multiple structures of øSimón, including the tail. Other areas worth exploring are the precise impacts of antibody titre and affinity on phage pharmacokinetics at different body sites, quantifying their consequence on therapeutic effects, and the influence of other components of the immune system, including phagocytes, on phage activity ^23,32,33^.

### Bacterial heteroresistance as a risk for treatment failure

Bacteria can use a vast repertoire of mechanisms to resist phage infection ^34,35^, and previous reports have demonstrated emergence of phage resistance during phage therapy ^36–39^. While this phenomenon can sometimes be beneficial, for example leading to collateral loss of antibiotic-resistance resulting in phage-antibiotic synergy ^37,39,40^, it can also be the cause for treatment adjustment ^41^, or failure ^38^. Therefore, we investigated if resistance to øSimón emerged during treatment, possibly contributing to the observed therapeutic failure.

We first assessed the ability and mechanism of the patient’s *B. bronchialis* to develop resistance to øSimón *in vitro*. After spotting øSimón on a lawn of wt *B. bronchialis*, we observed colonies growing in the middle of a lysis zone. We purified one of these colonies, confirmed it was phenotypically resistant to øSimón and sequenced its genome. Pairwise genomic comparison found a single mutation in this isolate, disrupting *btuB*, the gene coding for the outer membrane vitamin B12 receptor of the same name (Fig. S7a). The BtuB-deficient strain grew in the presence and absence of øSimón (Fig. S7b), had no fitness defect in minimal media (Fig. S7c-e), and an adsorption assay revealed that øSimón virions did not adsorb to this bacterial host (Fig. S7f). BtuB has been previously confirmed as either a sole, dual, or co-receptor for phages infecting the genera *Salmonella, Yersinia* and *Escherichia* ^42–45^. All of this suggests that øSimón may need BtuB for adsorption and that, as seen for many other phage-host pairs ^46^, loss of receptor function is a potential mechanism of *B. bronchialis* resistance to øSimón *in vitro*.

We then analysed the patient’s sputum samples over time for the presence of phage-resistant bacteria. Instead of starting our screening with individual colonies grown from each sputum sample, we took a population-wide approach. We adapted the inverted spot assay technique ^47^ whereby a high concentration of øSimón was used as a selection marker on semisolid media to detect putative phage-resistant *B. bronchialis*, with species confirmation performed using MALDI-ToF MS (Fig. 5a). This method allowed us to compare the number of *B. bronchialis* colonies that grew in the absence of øSimón (general population) against the number that grew in its presence (putative resistant population), thus calculating a ratio (Fig. 5b). Surprisingly, we discovered that before phage administration (day −1), a median of 2.3% (range: 1.7% to 11.6%) of the *B. bronchialis* population appeared to resist øSimón. This finding suggested baseline heteroresistance, defined as the presence of bacterial subpopulations with varied susceptibility to an antimicrobial (øSimón in this case), a phenomenon capable of contributing to treatment failure ^48^. To further investigate for heteroresistance, we randomly chose and purified 14 bacterial colonies from the pre-treatment sputum sample (day −1) and tested their efficiency of plating (EOP) for øSimón. A range of EOPs were identified across these pre-treatment colonies with most of them having values below 1, indicating reduced susceptibility to øSimón (Fig. 5c). These data supported the findings that the patient was infected with a heteroresistant population of *B. bronchialis*.

**Figure 5.**
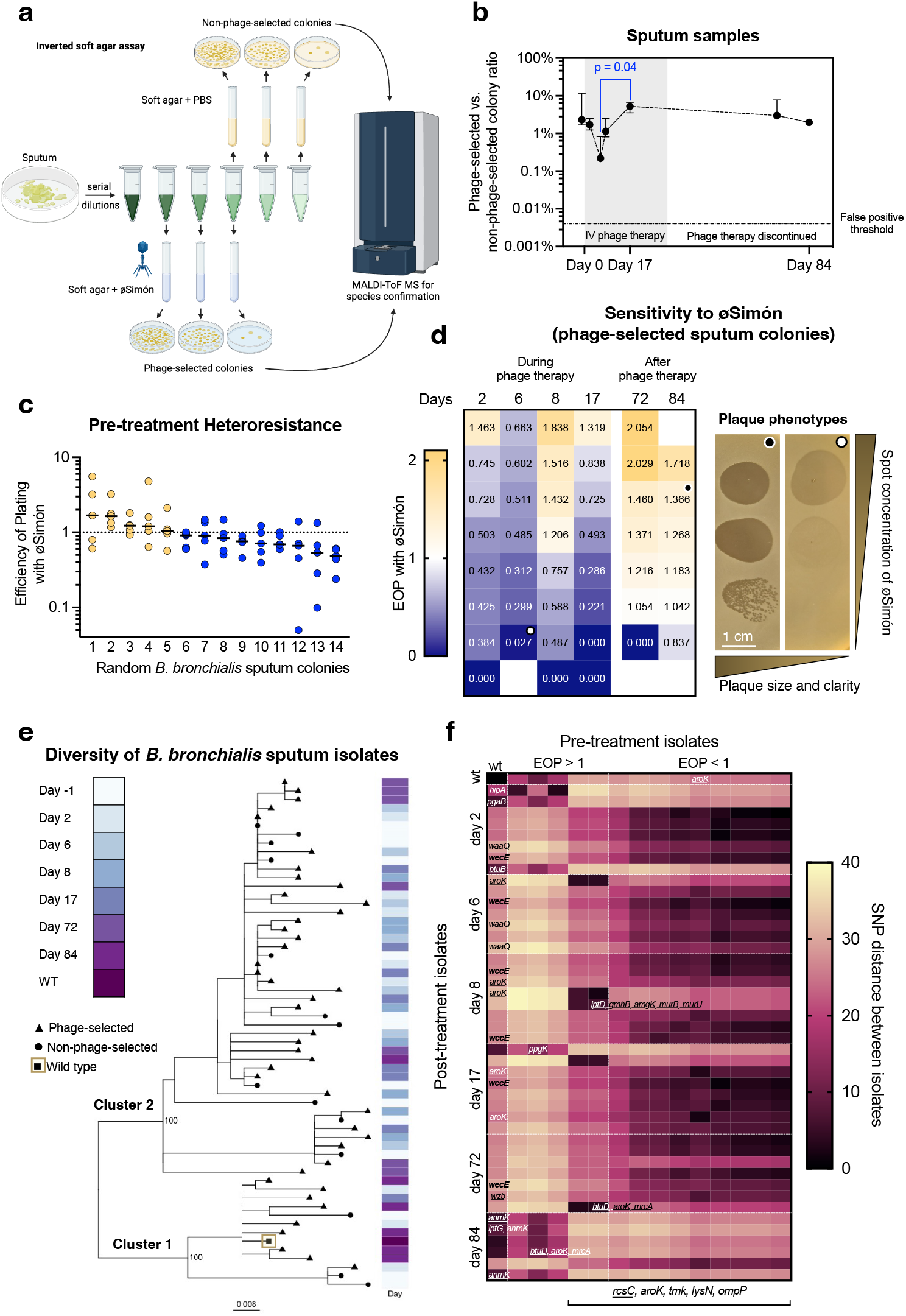
The patient was infected with a heteroresistant bacterial population. **a:** Phage-selection experiment. Serial dilutions of sputum were mixed with soft agar and either PBS (top) or øSimón (bottom), poured over hard agar plates and incubated. *B. bronchialis* colonies were identified through morphology and MALDI-ToF MS and counted. **b:** The ratio between non-phage-selected (PBS, top in panel a) and phage-selected (øSimón, bottom in panel a) was calculated for each sputum sample timepoint. Dotted line represents the median false positive threshold for the assay calculated on a clonal wt *B. bronchialis* culture (n = 3). Values are median ± 95% CI; n = 3; Kruskal-Wallis test, only p values < 0.05 plotted. **c:** Efficiency of plating (EOP) of øSimón on 14 random colonies from the pre-treatment sputum sample. All replicates plotted; bars are medians. EOPs > 1 in yellow, EOPs < 1 in blue. **d:** Heat map of the sensitivity/resistance spectrum to øSimón of sputum *B. bronchialis*. EOPs (n = 3; median plotted) of 44 phage-selected colonies from 6 different days during and after phage therapy. On the right, representative extremes of plaque size and clarity of øSimón on these strains. The white and black dots at the top right corner of each picture correspond to the strains used in the assay (marked with the same dots in the EOP panel). Scale bar = 1 cm. **e:** Maximum-likelihood tree of the 59 *B. bronchialis* colonies that were sequenced for this study. The tree depicts two clear clusters (100% node support values). **f:** SNP distance matrix between pre-treatment (columns), post-treatment (rows) and wt *B. bronchialis* (upper left corner). Columns ordered left-to-right in descending order of mean SNP distance to post-treatment isolates. Rows ordered by day of treatment and, within days, by descending EOP. The additional vertical dashed line in white separates pre-treatment isolates with EOPs higher and lower than 1, labelled respectively. Gene names in each cell represent mutations in a specific isolate from the rows, discovered after comparison against the corresponding isolate from the columns. The five genes under the heat map are mutated in all isolates of cluster 2, intact in cluster 1. Underlined genes: nonsense mutations or complete deletions; genes in bold: gain-of-function mutations; all the rest: missense mutations. All mutations are further detailed in Table S3 and the Supplementary Data File.

During and after treatment, the proportion of sputum colonies able to grow in the presence of phage remained relatively stable, but it reached a peak of 5.3% at day 17, coinciding with the bacteriological relapse seen in the patient (Fig. 2 and 5b). We randomly picked 6 to 8 colonies from the phage-selection plates (Fig. 5a) at each timepoint after initiating phage therapy (days 2, 6, 8, 17, 72 and 84, n = 44 colonies) and performed spotting and EOP assays with øSimón. In this subset, we also observed a range of EOPs, including five strains that were completely resistant to øSimón (EOP below the limit of detection of 10^−7^) (Fig. 5d). Mirroring the range in EOPs, we also observed variations in the size and/or clarity of the plaques øSimón produced on the isolates (Fig. 5d). Together, these findings revealed that before, during and after phage therapy, the patient harboured a bacterial population that was not uniformly sensitive to øSimón, and that instead of a sensitive/resistant dichotomy, the effectiveness of øSimón against the patient’s *B. bronchialis* fell within a spectrum. This solidified the hypothesis of heteroresistance and prompted an exploration of its genetic basis.

### Genetic analysis of bacterial heteroresistance

Using short-read whole genome sequencing of all 58 isolates (14 pre-treatment and 44 post-treatment) compared to the assembled genome of the wt *B. bronchialis* (host of isolation of øSimón, June 2022), we identified broad diversity, with two major clusters and several subclusters (Fig. 5e). Cluster 1 housed the wt strain, 3 of the 14 (18%) pre-treatment isolates, and 9 of the 44 (20%) post-treatment isolates. Cluster 2 was larger and included the remaining 11 pre-treatment and 35 post-treatment isolates. Using a SNP distance matrix (Fig. 5f) we noticed that the cluster differentiation corresponded to the EOP of the pre-treatment strains. Namely, all 3 strains belonging to cluster 1 had an EOP > 1 indicating high susceptibility to øSimón, whereas 9 of the 11 strains from cluster 2 had an EOP < 1. We then screened for specific genes and mutations that may explain the observed heteroresistance by comparing between the clusters in the pre-treatment bacterial strains. Strains from cluster 2 had a nonsense mutation in *rcsC*, which codes for a histidine kinase of the regulator of capsule synthesis (rcs) system. This system is activated upon cell surface stress, including outer membrane damage, lipopolysaccharide (LPS) synthesis defects, peptidoglycan perturbation and lipoprotein mislocalisation ^49^. It subsequently acts as a transcription regulator of associated genes, but also of those related to biofilm formation, motility and virulence ^50^. The system regulates the balance between innate and adaptive defence against phages ^51–53^, and mutations affecting it can confer partial phage-resistance ^54^. Cluster 2 also presented missense mutations in genes involved in DNA replication (thymidylate kinase [*tmk*]), amino acid metabolism (shikimate kinase [*aroK*]; 2-aminoadipate transaminase [*lysN*]), and the outer membrane porin *ompP*, although their contribution to decreased susceptibility to øSimón is uncertain.

Finally, we looked for mutations in the post-treatment, phage-selected isolates and found that compared to the wt *B. bronchialis* strain, there was a median of 27 SNPs per strain (range 4 – 32), with no correlation between the number of SNPs and susceptibility to øSimón (Fig. S8 and Supplementary data file). Notably, a bacterial isolate recovered from sputum after 2 days of phage therapy had a total deletion of *btuB*, the gene that encodes for the vitamin B12 transporter that we showed was important for øSimón-resistance *in vitro* (Fig. S7). This isolate was completely resistant to øSimón. Another isolate from day 2 had a missense mutation in the toxin gene *hipA*, which is part of a known toxin-antitoxin phage defence system ^55^. Additionally, we found that many of the phage-selected strains, regardless of their genetic background (cluster 1 or 2), also harboured mutations in cell envelope-associated genes, including cell wall synthesis (*anmK, mrcA, amgK, murB, murU*), LPS biosynthesis (*pgaB, ppgK, waaQ, wecE, gmhB*), LPS export (*lptD, lptG*), and capsular and extracellular polysaccharide synthesis (*wzb*) (Fig. 5f, Table S3), all potentially impacting their susceptibility to øSimón to varying degrees and through different pathways ^56–60^. Taken together, these data show that our patient was infected with genetic variants of *B. bronchialis* that had a diverse spectrum of susceptibility to øSimón, even before phage treatment started. After treatment initiation, strains with reduced susceptibility to øSimón persisted, with several defined mutations, including loss of the putative receptor *btuB*, likely explaining the phenotype.

The pre-treatment phage heteroresistance we identified likely contributed to the failure of phage therapy. However, our findings should be interpreted with nuance. First, the proportion of bacteria with reduced sensitivity to øSimón did not dramatically increase during phage administration (Fig. 5b), suggesting these subpopulations did not possess a significant advantage under phage selection *in vivo*. Second, the isolates obtained during and after phage treatment were distributed across clusters 1 and 2 at a similar ratio as the pre-treatment isolates. This could indicate that the observed genomic changes reflect natural pathogen variability or in-host evolution, rather than direct selection by phage treatment. In any case, we argue that testing for heteroresistance prior to phage treatment would be advisable in future patients.

### Clinical implications

The resurgence of phage therapy in the context of challenging AMR bacterial infections has highlighted its many remaining unknowns. Here, we provide a detailed analysis of a clinical case of phage therapeutic failure and uncovered important insights with clinical implications for the future care of phage therapy patients. Using longitudinal clinical samples, we first identified the complex dynamics that are taking place between phages, bacteria and human immune responses. When administered intravenously, phage was able to reach the patient’s sputum, however phage activity was short-lived. Two main factors appeared to be at play: pre-existing antiphage immunity and bacterial heteroresistance to phage. Importantly, we have shown that in the setting of a chronic bacterial infection, bacterial prophages could act as host immune stimulators, leading to pre-treatment cross-reactive antibodies that can target the therapeutic phage. This pre-treatment phage neutralisation capacity, combined with subpopulations of infecting bacteria with reduced susceptibility to the phage, create a recipe for clinical failure that has been under-appreciated.

We recommend screening a patient’s serum for antiphage immunity (serum neutralisation) well before undertaking clinical production of a specific phage. The current Phage Australia clinical protocol suggests testing for antiphage antibodies during and after phage therapy ^12^. We suggest that, in the presence of alternative phages suitable for therapy, those that are susceptible to pre-existing immunity should be considered lower priority. In the absence of an alternative phage for a patient requiring urgent treatment, however, pre-existing immunity should not be considered a contraindication. Second, we recommend choosing a phage or combination of phages for clinical use by testing them against a patient’s bacterial population instead of a single colony. Homogenous colony morphology or growth kinetics should not be used as evidence to rule out possible bacterial heteroresistance. Instead, the inverted top agar assay described in this study could be used for that purpose in a cost-effective manner. An increasing amount of evidence recognises that, especially in chronic infections, patients can harbour heterogenous populations of a pathogen ^61–63^, and a single colony might not be representative of the population’s susceptibility to phage therapy. These clinical recommendations are easily applicable to all prospective phage therapy patients but are particularly relevant in patients with recurrent, chronic or complex multisite infections. Likewise, it will be important for those planning clinical trials to consider these variables in trial design and interpretation, including the collection of appropriate samples throughout the study. We hope that as global efforts to bring phage therapy into widespread clinical practice continue, careful analysis of clinical cases and results from well-designed clinical trials will keep providing useful and actionable insights to maximise the effectiveness of this promising antimicrobial strategy.

## Materials and Methods

### Bacterial growth, quantification, and storage

*B. bronchialis* was grown on heart infusion (HI) broth (Oxoid, CM1032), supplemented with 1.5% agar when necessary, aerobically, with constant shaking for liquid cultures, at 37°C for 48 h. Bacterial quantification was performed using the spread-plating technique, with at least two successive 10-fold serial dilutions of the relevant sample yielding countable colonies. When needed, bacterial cultures were diluted 2:3 in 80% glycerol for storage at −80°C.

### Phage isolation, purification, quantification and efficiency of plating (EOP)

Phage øSimón was isolated from raw sewage samples (Melbourne, VIC, Australia, May 2022). Aliquots of 100 ml of centrifugated and filtered sewage were combined with 1 ml of a turbid *B. bronchialis* liquid culture and supplemented with 10× HI, and 10 mM each of CaCl_2_ and MgSO_4_. Mixtures were incubated as above, the resulting lysates were purified with the Phage-on-Tap protocol ^64^, with or without buffer replacement, for general lab stocks. Phage suspensions needed for Immunology work underwent further purification as performed in ^24^. All phage products were stored at 4 °C. Phage quantification and EOP testing were performed with a slightly modified double overlay agar assay ^65^ using 4 ml of molten soft (0.4%) HI agar and 1 ml of a *B. bronchialis* culture at an optical density (OD_600_) of 0.3 (previously tested to correspond to a bacterial density of 8 × 10^8^ cfu/ml), per plate. Plates were always read after 48 h of incubation, and the average number of plaques in at least two plates was recorded and compared as a percentage of the plaques formed on the original host of isolation under the same conditions.

### Bacterial and phage kinetics

Growth kinetics were determined using a 96-well plate format and spectrophotometry (BioTek Epoch 2), measuring OD_600_ every 15 min with a starting inoculum of 10^6^ cfu/well, and phage pfu at multiplicities of infection (MOI) of either 0, 0.01, 0.1, or 1. For experiments comparing bactericidal activity in liquid and sputum, cocultures were set up in 10 ml volumes, at a starting bacterial density of 10^7^ cfu/ml and MOIs of either 0, 0.01, or 0.1. Artificial sputum medium was commercially acquired (Biochemazone, Canada, BZ316), handled, stored and reconstituted according to the manufacturer’s instructions.

### Phage one-step kill curve and adsorption assay

Bacteria from liquid cultures and phage from a purified stock were mixed at an MOI of 0.01 (10^8^ cfu/ml) in HI broth, with cation supplementation as above. The assay was run in 10 ml total volumes. The suspensions were incubated, and at each timepoint, 1 ml samples were taken and filtered with a 0.2 μm syringe filter (Pall, 4612). This filtrate was serially diluted in PBS and plated in duplicate for quantification of free phage particles. A control with no bacteria was routinely run in adsorption assays.

### Transmission electron microscopy (TEM) and immuno-TEM

Droplets of phage suspension (10 μl) were placed on copper TEM grids (200 mesh; SPI) with carbon-coated ultrathin formvar film. They were left for 30 s and then dried using filter paper. For negative staining, the grids (face side down) were put to droplets (10 μl) of uranyl acetate water solution (1% w/v) and incubated for 30 s. Finally, the grids were dried with filter paper and examined under a transmission electron microscope (JEM-1400 Plus) at an accelerating voltage of 80 kV. In the induced prophage suspension, imaging was hindered by contamination and low virion concentration. For immuno-TEM experiments, the IgG fractions of the patient’s serum samples were first purified using protein G magnetic beads (New England Biolabs, S1430S) largely according to the manufacturer’s instructions. Briefly, after washing beads in binding buffer (0.1 M sodium carbonate, pH = 8.0) three times, serum diluted 1:3 with binding buffer was added to beads and then incubated with rolling and tilting for 15 min at room temperature. Beads were placed on a magnetic stand for 30 s, the depleted serum removed and then the beads washed four times with 1 ml binding buffer. Following the washes, IgG was eluted in 0.2 M glycine pH = 2.5 and transferred to a tube containing a neutralising volume of 1 M Tris-HCl pH = 9.0. The sample was diluted 1:4 in PBS, filtered (0.2 μm) and concentrated sequentially through 10 kDa Amicon Ultra Centrifugal filters (Merck Millipore, UFC8010 and UFC5010). The A280 of the concentrated, purified IgG was measured by NanoDrop (NanoDrop Technologies LLC) and a 0.1% Ce of 1.36 applied to determine the IgG concentration (at least 0.2 mg/ml for immuno-TEM). Phages were left to adsorb to the TEM grids for 10 min, followed by three 1-min washes on droplets of 1% bovine serum albumin (BSA) in PBS. Grids were then incubated for 2 h on a mixture containing 10 μl of 1% BSA in PBS and 5 μl of patient-derived IgG. After three additional 1-min washes, grids were incubated for 1 h on a mixture of 10 μl of 1% BSA in PBS and 5 μl of gold-conjugated goat-anti human IgG antibodies (RRID: AB_2337747) (diluted 1:10 in PBS). This was followed by three washes with PBS, and standard negative staining and examination.

### Phage DNA extraction, sequencing and bioinformatics

One ml volumes of high-titre phage lysate (> 10^8^ pfu/ml whenever possible) were treated with DNase (1 mg/ml) and RNase A (12.5 mg/ml) for 2 h at 37 °C, followed by an inactivation step of 5 min at 75 °C. We then used the Norgen phage DNA kit (Norgen Biotek Corp., Thorold, ON, Canada) to extract DNA following the manufacturer’s instructions. DNA concentration was measured using a Qubit® Fluorometer, quality confirmed with the 260/280 nm absorbance ratio, and absence of bacterial contamination was confirmed by 16S rRNA PCR and gel visualisation of the DNA samples. Phage øSimón was sequenced on an Illumina HiSeq platform, and with Oxford Nanopore Technologies (ONT) long-read sequencing with a GridION device on FLO-MIN114 R10.4.1 flow cells. The reads were trimmed using trimmomatic (v0.39) ^66^, assembled with Unicycler (v0.5.0) ^67^, and annotated using Prokka (v.1.14.6), the RAST server and the PHROG database ^68–70^. The genome was set to start at the large terminase subunit (*terL*) gene ^71^, polished with Pilon (v.1.22) ^72^. manually screened for lysogeny indicators and assessed for AMR determinants using AMRFinderPlus (v.3.11.26) ^73^ and Abricate (v.1.0.1) (https://github.com/tseemann/abricate). Phylogeny was investigated using vContact2 ^74^ and the gene voting approach ^75^, both using the Inphared database (version 1Dec2021) ^76^. Gene similarity between øSimón and *B. bronchialis* prophages was studied using blastp (v.2.15.0+) and genome maps were visualised using the R package *gggenomes* (v.1.0.0).

### Production of clinical-grade phage øSimón

We followed a slightly modified version of a previously used protocol ^10^. The phage was amplified in 1,000 ml HI broth. The lysate was centrifugated for 30 min at 4,000 × g to remove larger bacterial debris, and then filtered through a depth filter from 5 μm to 0.2 μm, monitoring the pressure not to exceed and compromise the filter specifications (<2 bar). The lysate was passed once again through a sterilisation-grade (0.2 μm) filter and transferred to a bacteria-free production facility. The lysate was diluted 1:10 in PBS and flushed through an ÄKTA Flux tangential flow filtration (TFF) system (Cytiva) equipped with a 300 kDa microfiltration hollow fibre cartridge. This process was repeated until the HI broth was completely replaced by PBS. The lysate was filter-sterilised again, and the phage titre measured. Endotoxin removal was performed using EndoTrap®HD affinity columns, and the endotoxin concentration was assessed with Endosafe® PTS™ cartridge (Charles River laboratories, MA, USA) kits. After achieving a satisfactory endotoxin level, we processed the samples under strict sterile conditions, packed them in sealed glass vials with syringe access (Medisca, 9497-01), and sent them to Eurofins BioPharma Product Testing in Sydney, Australia for third-party validation for sterility (USP71) and endotoxin (USP85).

### Handling and storage of clinical specimens

Serum samples were aliquoted into 200 μl volumes and kept at −20 °C, minimising freeze/thaw cycles. For sputum samples, we first measured their volume and mixed them 1:1 with a 1:10 solution of dithiothreitol (Sputolysin®, CAS 578517) in sterile water with vortexing for 1 min. Bacterial and phage quantification were performed at this stage. For long-term storage for other experiments, each sample was mixed 1:1 with sterile glycerol broth (60 ml glycerol, 2 g tryptone, 1 g yeast extract, 2 g NaCl, in 200 ml deionised water) in cryovials and stored at −80 °C.

### Serum and sputum neutralisation assays and calculation of the phage inactivation rate (*k*)

The clinical samples were diluted 1:10 in PBS, filter-sterilised (0.2 μm syringe filter), and then further diluted in PBS in 10-fold serial dilutions. Working volumes of 1 ml of these dilutions, and a PBS-only control, were spiked with 10^6^ pfu each of phages øSimón and øFaith (unpublished phage, isolated in our lab against *Pseudomonas aeruginosa* strains from patients with cystic fibrosis). The samples were incubated at 37 °C for 30 min with constant shaking. At the endpoint, phage quantification using the double overlay agar assay was performed for each phage on their respective hosts of isolation. Neutralisation was calculated as the percentage of active phage at the endpoint compared to the initial phage inoculum for each condition. The phage inactivation rate (*k*) was calculated with the formula:

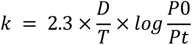

Where D is the reciprocal of the serum or sputum dilution in PBS, T is the time of incubation, and P0 and Pt are the phage titres at the beginning and endpoint of the assay, respectively ^19,21^.

### Indirect ELISA and cross-reactive ELISA

We based our protocol in ^20^ with some modifications. Individual wells of MICROLON™ high binding 96-well plates (Greiner, 655061) were coated with 100 μl of either pure coating buffer (carbonate-bicarbonate with pH = 9.6, Sigma C3041), phage øSimón calculated at 10^5^ pfu/well, induced prophage at 10^3^ virions/well, or phage vehicle (obtained after filtering phage solutions with a 0.02 μm syringe filter, Whatman 68091002); the latter three diluted in coating buffer. Plates were covered with foil and incubated at 4 °C for 20 h. The wells were emptied, washed 4 times with 300 μl of PBS + 0.05% Tween 80 (PBST), blocked with 200 μl of 3% skim milk powder in PBS, and the plates incubated as above. The blocking solution was shaken off, and the plates patted dry with paper towels. A further blocking step was performed with 3% bovine serum albumin (BSA) in PBS at 37 °C for 30 min, before washing with PBST. The patient samples (serum or filter-sterilised sputum) were diluted in 3% BSA in PBS, added to the respective wells in volumes of 50 μl, incubated at 37 °C for 1 h, and washed with PBST. Next, volumes of 50 μl of 1:500 goat anti-human IgA + IgG + IgM-AP (alkaline phosphatase)-conjugated antibody (RRID: AB_2337605) were added and the plates again incubated and later washed with PBST. Finally, the p-Nitrophenyl phosphate substrate (Sigmafast^TM^, N2770) was added in a volume of 50 μl and the reaction left to develop for 20 min. The assay was stopped by adding 50 μl of 2 M NaOH and the plates were read with a spectrophotometer at 450 nm. For the antibody cross-reactivity testing the same protocol was followed except for a “pre-capture” step involving the passage of individual serum samples by wells covered with øSimón, prophage, or phage vehicle in coating buffer, for 1 h before transferring them into new plates and wells for a regular ELISA as previously described.

### Prophage induction

A stationary phase *B. bronchialis* culture was diluted 1:10 in HI and left for incubation for 6 h. The culture was split into two 10 ml aliquots, the first one received a PBS spike (spontaneous induction control), and the second one a mitomycin C at 0.3 mg/ml spike, in equal volumes. Both cultures were left for overnight incubation and purified as above ^24^. Prophage induction was confirmed through TEM and phage DNA sequencing.

### Inverted soft agar assay for phage-selection

Sputum samples were 10-fold serially diluted in PBS. Then, 100 μl of the dilutions were mixed with 4 ml of molten soft (0.4%) HI agar and 100 μl of either PBS (no phage selection) or a stock of øSimón for a concentration of 10^7^ pfu/plate (phage selection), before being poured over plates containing a first layer of standard HI agar. Plates were left for 48 h of incubation and, for counting, *B. bronchialis* colonies on each plate were identified through visual assessment of their morphology, followed by species confirmation using MALDI-ToF MS (Bruker, Biotyper microflex®). Normal microbiota from the upper respiratory tract was consistently seen at concentrations 2 log lower than *B. bronchialis* and thus did not represent a significant confounder in this process. Phage-selected and non-phage-selected colonies underwent two rounds of single-colony purification before being grown in liquid culture and stored as above.

### Bacterial DNA extraction and sequencing

Genomic DNA was extracted from *B. bronchialis* cultures using the GenFind V3 Reagent Kit (Beckman Coulter) as per manufacturer’s instructions. Illumina libraries were prepared using the Illumina DNA Library Prep Kit, (M) Tagmentaton (Illumina) and sequenced on the Illumina NovaSeq 6000 platform to produce paired-end reads. The wt *B. bronchialis* isolate also underwent long-read Nanopore sequencing with a GridION device on FLO-MIN114 R10.4.1 flow cells (Oxford Nanopore Technologies). Sequencing libraries were barcoded with the native barcoding kit SQK-RBK114.96 in accordance with the protocol RBK_9176_v114_revM_27Nov2022.

### Bacterial genomics

A *de novo* assembly of the genome of wt *B. bronchialis* was first constructed using Trycycler (v0.5.4) with long read data ^77^, polished with matching short reads using Polypolish (v0.5.0) ^78^ and annotated with Bakta (v1.7) ^79^. The short-read data from the other isolates also underwent quality control (https://github.com/s-andrews/FastQC) before the genomes were assembled with Unicycler (v0.5.0) ^67^ and annotated with Prokka (v.1.14.6) ^68^. We identified SNPs between the wt *B. bronchialis* reference genome and other sputum isolates using the Nextflow implementation of the Reddog pipeline v1beta.11 using default parameters (https://github.com/scwatts/reddog-nf). The resulting SNP alignment consisted of 250 sites. Complementarily, the wt genome was manually parsed for insertion sequences, whose nucleotide sequences were then mapped against the short reads from other isolates using ISMapper v2.0.0 ^80^, to detect alternative gene disruptions in these isolates. To identify the absence of whole genes in pairwise comparisons between isolates we used panaroo (v1.2.4) ^81^ using thresholds of ≥95% for gene coverage and ≥5x for depth. Phylogeny was studied by constructing a maximum-likelihood tree from the SNP alignment obtained above, with the GTR+Γ model, using RAxML (v8.2.9) ^82^ with 1,000 bootstrap replicates, then visualised with “ape” (v5.7.1) ^83^ and “ggtree” (v3.6.2) ^84^ in R. The pairwise SNP distance matrix was created using “harrietr” (https://github.com/andersgs/harrietr).

### Graphing, statistics, and reproducibility

Graphing and statistical analyses were performed with GraphPad Prism 9 (GraphPad Software, Inc.). Figures 4d and 5a were crafted using BioRender. All *in vitro* experiments were performed in triplicate, unless specified otherwise on the respective figure legends, with at least two technical replicates each. Where appropriate, due to the small number of replicates, medians were presented. To decide between the use of parametric or nonparametric statistical analyses, the data were tested for normality using the Shapiro-Wilk method. The threshold value of two-tailed p < 0.05 was used for statistical significance. Observational results, including those from TEM, immuno-TEM, and phage plaque morphology are representative of at least 100 observations each.

## Data availability

All bacterial and phage genomes, sequencing data, and metadata derived from this study have been deposited on NCBI under the BioProject number PRJNA1180749 (Table S4).

## Ethical statement

All procedures performed in this study were in accordance with the ethical standards of the institutional research committee (Alfred Hospital Ethics Committee approval number 230/24) and with the Helsinki Declaration (as revised in 2013). Written informed consent to receive phage therapy was obtained from the patient. Data from this clinical case forms part of the STAMP Study ^12^, trial registration number (registered on ANZCTR 10 Nov 2021): ACTRN12621001526864; WHO Universal Trial Number: U1111-1269-6000.

## Acknowledgements

We thank Dr Ryan Burrows (University of Melbourne & Melbourne Water) for his help establishing the collaboration that granted us access to the raw sewage samples used to isolate øSimón. Thanks to Christina Rootes, Dr Christopher Pace, and Dr Duen Siauw for useful feedback on the structure and readability of the manuscript, and to Ruzeen Patwa for her help in logistics and laboratory management. Thanks to Simón Usúcar Gordillo, after whom phage øSimón was named (*que tu vida esté llena de éxito y alegría, tu tío te ama*). The Monash Antibody Discovery Platform is supported by funding from Bioplatforms Australia (enabled by NCRIS). This work was supported by Monash eResearch capabilities, including M3, and the Alfred Hospital and Monash University funding of the Victorian Bacteriophage Therapy Program (VIC*Phage*) and the Monash Phage Foundry. Jeremy J. Barr and Anton Y. Peleg also acknowledge the Australian National Health and Medical Research Council (NHMRC) Investigator L1 Grant (2023/GNT2026130) and Practitioner Fellowship (APP1117940), respectively, and the Frontier Health Medical Research support (RFRHPI000017).

## Author contributions

Conceptualisation: FGA, JJB, AYP.

Laboratory experimental work: FGA, DS, MBu, DMP, MP, DKo, MJR, KP, JW, HR.

Genomics and bioinformatics: DS, MBe, MBu, SD, JH.

Clinical monitoring and follow up: FGA, SK, BJG, YH, DKe, TK, AYP.

Data curation, analysis and visualisation: FGA, AYP.

Provision of resources: CR, JJB, AYP.

Original preparation of the manuscript: FGA, AYP.

All authors reviewed, edited and approved the manuscript.

