## Supplementary Figures 1-8, Tables 1-3 and Notes 1-3 for "Mechanistic insights and clinical implications of cross-reactive anti-prophage antibodies and bacterial heteroresistance on phage therapeutic failure"

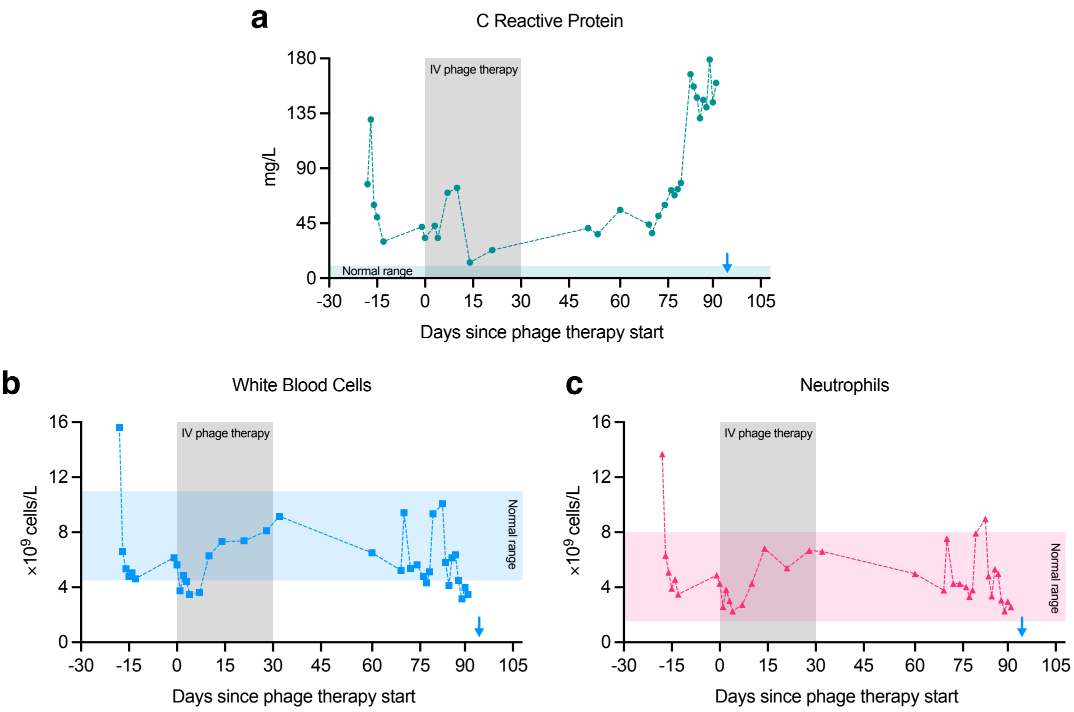


**Supplementary Figure 1. Immune and inflammatory markers during phage therapy.** Levels of C-reactive protein (**a**), white blood cells (**b**), and neutrophils (**c**) in the patient over the course of the clinical history. Grey shaded zones in each panel represent the 30-day period of phage administration, whereas the coloured shaded zones represent the normal ranges for each marker. Blue arrows demark the day of patient death.

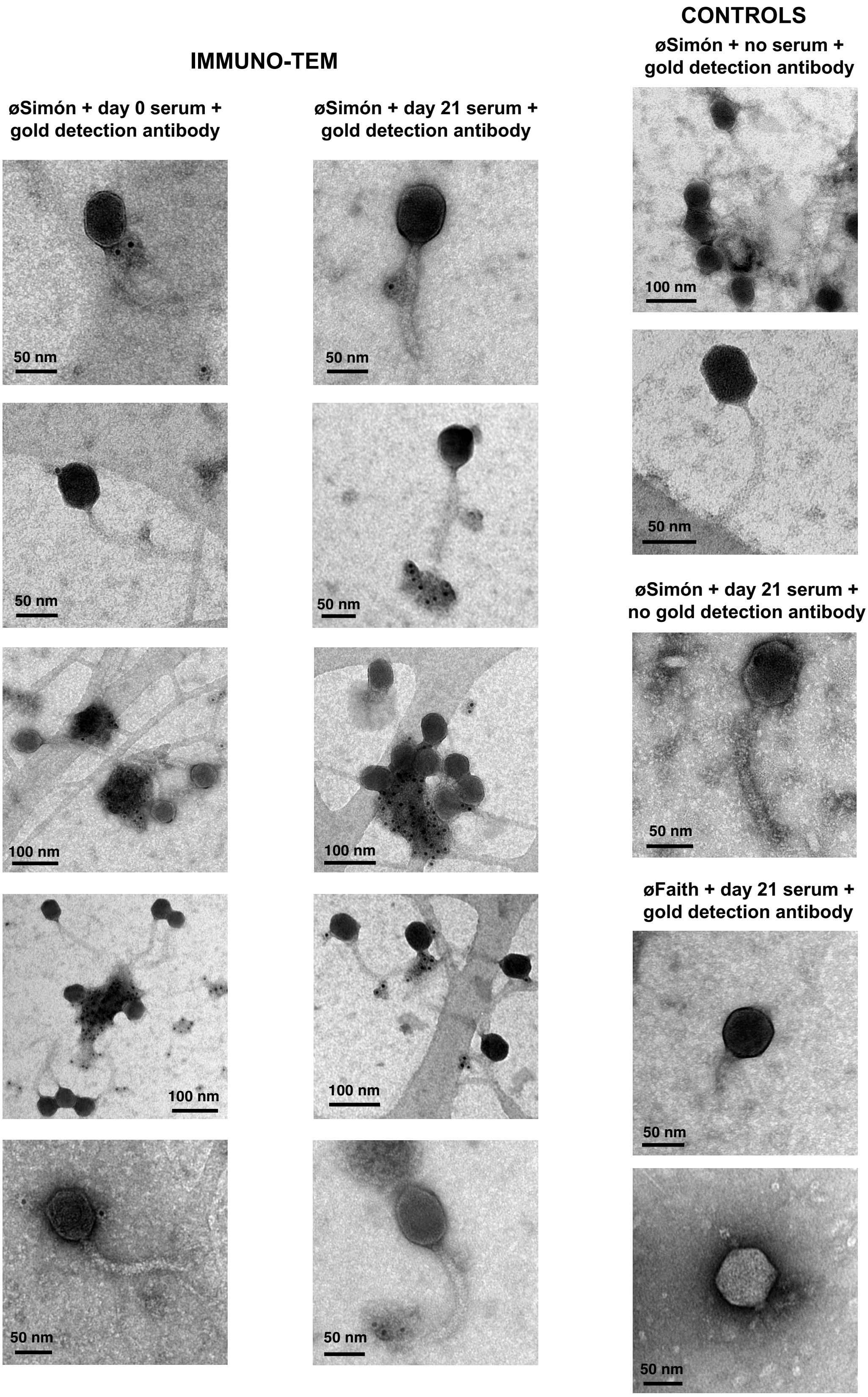


**Supplementary Figure 2. Immuno-TEM (Transmission Electron Microscopy).** Additional micrographs of phage øSimón incubated with patient's serum from before phage therapy (Day 0) or after 21 days of phage therapy, and a gold-labelled anti-human IgG detection antibody (black dots) (left and middle columns, respectively). Assorted controls for the experiment on the right column. Scale bars depicted at the bottom corner of every micrograph.

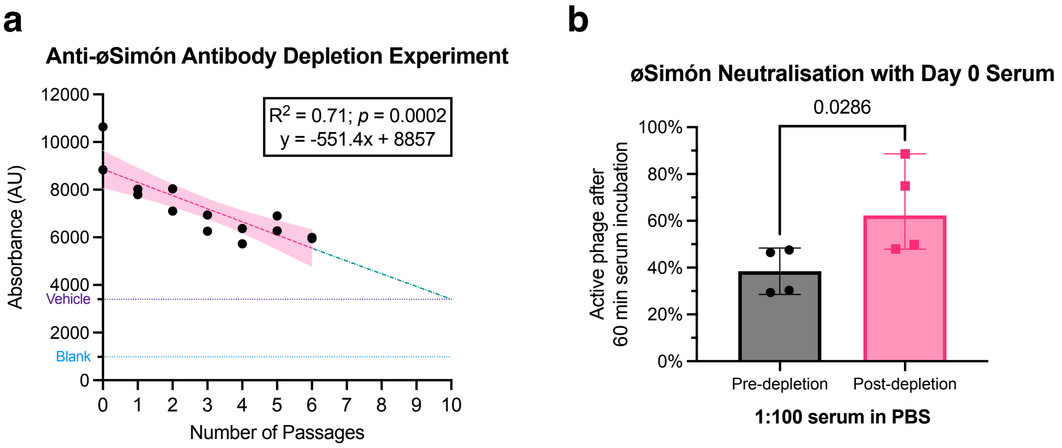


**Supplementary Figure 3. Depletion of anti-øSimón antibodies reverses the neutralisation effect of the patient's serum. a:** Correlation between the number of successive passages of serum on øSimón-coated wells, and the ELISA signal for anti-øSimón antibodies. Simple linear regression with 95% CI shaded zones depicted in magenta. Using the equation from the regression, it was predicted that the signal for anti-øSimón antibodies would reach the level of the vehicle control (purple line) after 10 passages. **b:** Neutralisation assay of a $10^6$ pfu/ml dose of øSimón after 60 min of incubation in day 0 patient serum 1:100 in PBS (black circles), or the same serum sample after 12 passages for anti-øSimón antibody depletion (magenta squares). The values for the post-depletion column were corrected by subtracting the amount of residual øSimón from the passages used for antibody depletion. Bars are medians ± 95% CI; Mann-Whitney test; two-tailed.

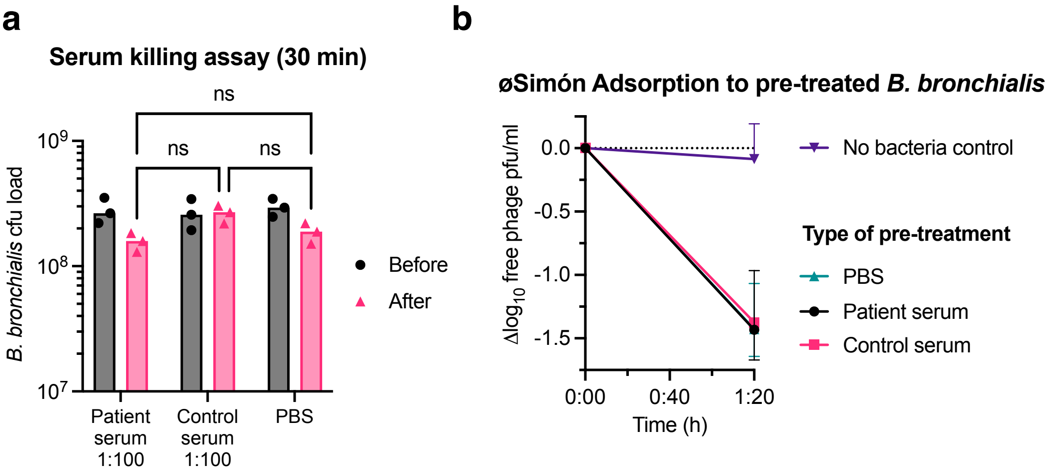


**Supplementary Figure 4. Patient serum does not inhibit øSimón through effects on the bacterial surface. a:** Bactericidal effect of patient serum, control serum or a PBS control over a *B. bronchialis* population. Bacteria were counted before (black circles) and after (magenta triangles) 30 minutes of treatment (bars are medians; n = 3; two-tailed two-way ANOVA: ns: not significant). **b:** The bacterial populations from panel (a) were used in an adsorption assay with øSimón. There were no differences in øSimón's ability to adsorb to any of these bacterial cells after 80 minutes of coincubation (median ± 95% CI; n = 3)

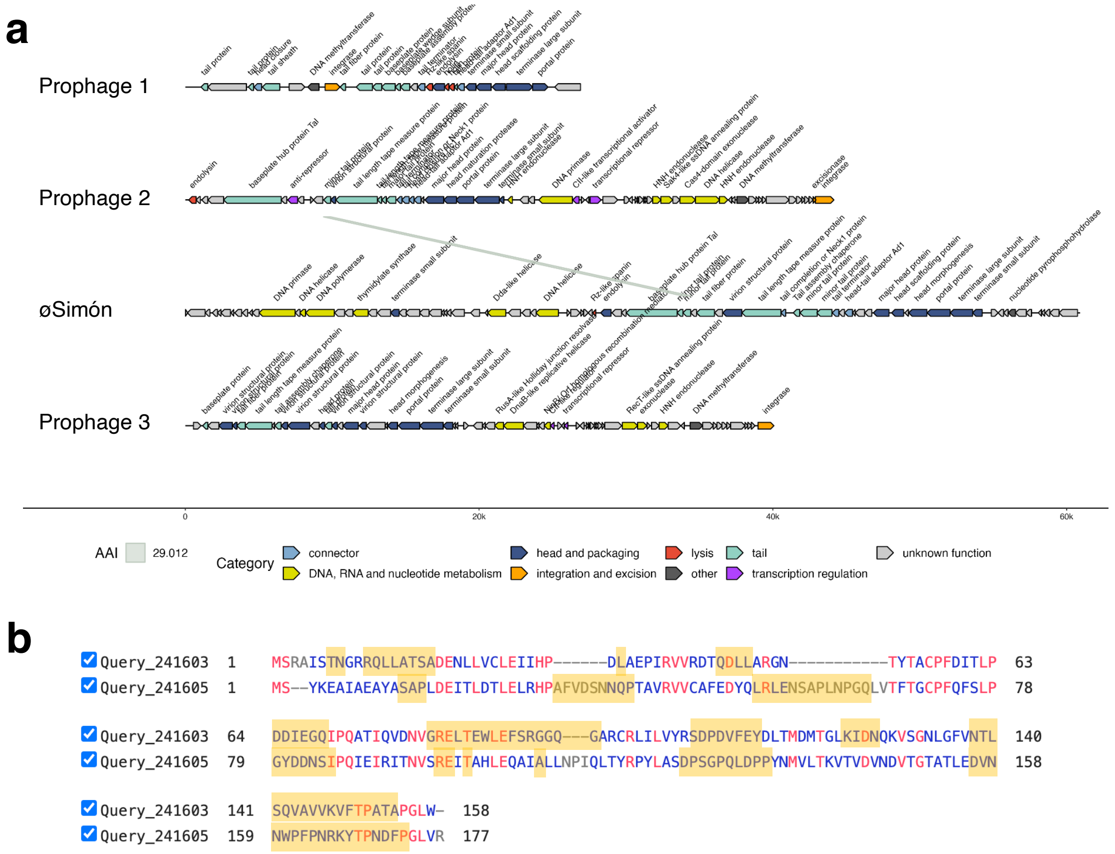


**Supplementary Figure 5. Possible origin of the cross-reactivity between øSimón and *B. bronchialis* prophages. a:** Assembled and annotated genomes of the three prophage regions found in the genome of wt *B. bronchialis,* and phage øSimón. Genes are colour-coded according to the predicted function. Modest amino acid identity (AAI, 29%) was found between minor tail protein genes of prophage 2 and øSimón (> 85% query cover). No other significant homologies were found. **b:** Aligned amino acid sequences of the minor tail protein of prophage 2 (top) and øSimón (bottom). Residues in red are identical in both proteins. Yellow boxes highlight areas predicted to be B-cell recognisable epitopes [1].

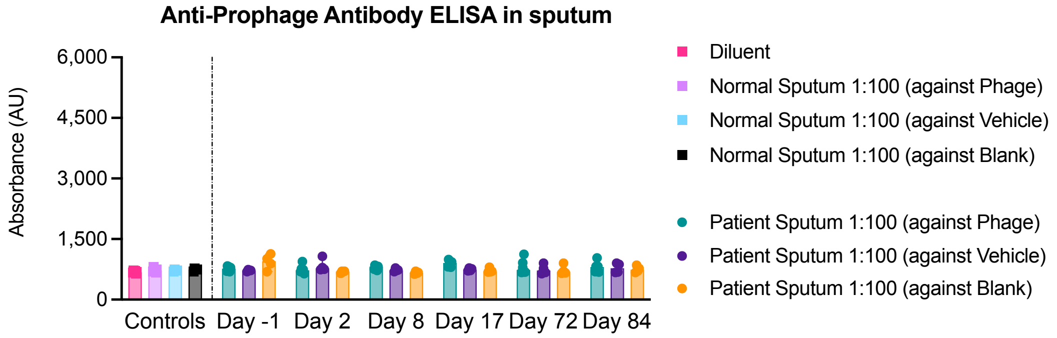


**Supplementary Figure 6.** Indirect ELISA testing to detect antibodies against *B. bronchialis* prophages in the patient's sputum from 6 different days. Control sputum came from a patient with cystic fibrosis. Vehicle control was obtained after filtering phages out using a 0.02 μm filter, blank control was PBS. Bars are median ± 95% CI; all replicates from two experiments plotted; Kruskal-Wallis test, all comparisons were not significant.

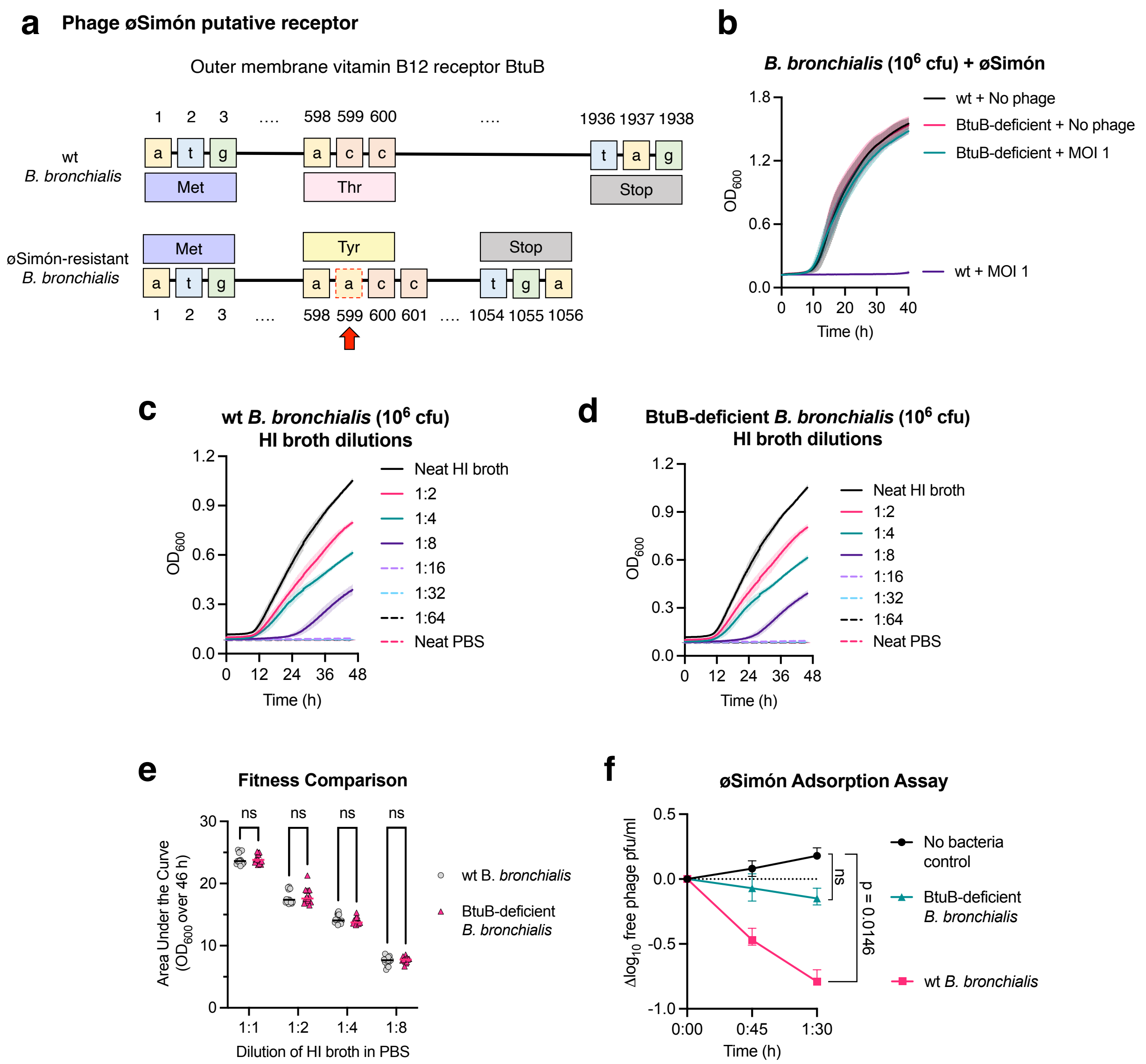


**Supplementary Figure 7.** ***In vitro*** **mechanism of resistance against phage øSimón. a:** Representation of the *btuB* gene sequence in wt *B. bronchialis* (top) and a mutant resistant to øSimón (bottom). The red arrow and box indicate the insertion of an alanine at position 599, leading to a premature stop codon in the phage-resistant bacterium. **b:** Bacterial growth curves of wt *B. bronchialis* and the BtuB-deficient variant presented in panel (a), with and without phage (mean ± SD; n = 3). **c and d:** Bacterial growth curves of wt and BtuB-deficient *B. bronchialis,* respectively, in serial 1:2 dilutions of heart infusion (HI) broth in PBS (mean ± SD; n = 3). **e:** For the dilutions from panels (c) and (d) that supported *B. bronchialis* growth*,* comparison of the area under the curve between wt (grey) and BtuB-deficient (pink) isolates (bars are medians; all technical replicates from three experiments plotted; 2-way ANOVA; two-tailed). ns: not significant. **f:** Adsorption assay of øSimón to wt or BtuB-deficient *B. bronchialis* over 90 minutes (median ± 95% CI; n = 3; Kruskal-Wallis test).

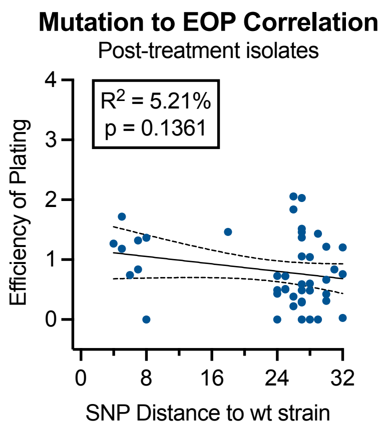


**Supplementary Figure 8.** Linear correlation between EOP and SNP distance to wt strain in the post-treatment isolates (n = 44) shows a negligible association. Dashed lines represent the 95% confidence intervals.

**Supplementary Table 1: Antimicrobial susceptibility testing (MIC reported in μg/ml) of the patient's *Bordetella bronchialis* isolates, demonstrating the development of resistance to multiple agents over a 7-year period**

| Antimicrobial | Patient's age | | |
|---|---|---|---|
| | Age 22 | Age 20 | Age 15^ |
| Piperacillin-tazobactam | >256 | >256 | 4 |
| Ceftriaxone | | >32 | |
| Ceftazidime | >256 | >256 | 8 |
| Ceftazidime-avibactam | >256 | >256 | |
| Ceftolozane-tazobactam | >256 | | >256 |
| Cefepime | >256 | | |
| Cefiderocol | No zone | | |
| Imipenem | >32 | >32 | >32 |
| Imipenem-relibactam | 4 | | |
| Meropenem | >32 | >32 | |
| Meropenem-vaborbactam | 4 | | |
| Aztreonam | >256 | >256 | R* |
| Gentamicin | | | >256 |
| Tobramycin | | | R* |
| Amikacin | >32 | | R* |
| Cotrimoxazole | >32 | 0.5 / 1 | R* |
| Ciprofloxacin | >32 | >32 | 16 |
| Moxifloxacin | >32 | >32 | |
| Doxycycline | 2 | 1 | 1 |
| Minocycline | 2 | 1 | 0.5 |
| Tigecycline | 0.25 | 0.5 | 0.25 |
| Eravacycline | 0.125 | | |
| Erythromycin | | >256 | |
| Clarithromycin | | | >256 |
| Azithromycin | >256 | | >256 |
| Fosfomycin | 8 | | 32 |
| Colistin | >64[#] | | 16 |

^historic data, isolates no longer available; *no MIC data available; #tested 3 times across the year, with one discrepant result of 2. Methodology available in Supplementary Note 1.

**Supplementary Table 2. Information on batches of øSimón clinical-grade preparations**

| # | Titre at packaging ($\log_{10}$ pfu/ml) | Date of packaging | Administered clinically | Notes |
|---|---|---|---|---|
| 1 | 7.73 | 06-12-2022 | Yes, fully | 33 vials, used for therapy from days 1 to 16, finishing on 07-01-2023 |
| 2 | 7.00 | 15-12-2022 | Yes, partially | 85 vials, used to complete the phage therapy protocol (days 17 to 30), finishing on 19-01-2023 |
| 3 | 7.00 | 17-02-2023 | No | 132 vials |

**Supplementary Table 3. Additional information on relevant genes found to be mutated in cluster 2 (*) and phage-selected *B. bronchialis* isolates.**

| Symbol | Name | Function / description | Length (AA) | Mutation Consequence(s) | Important References |
|---|---|---|---|---|---|
| *rcsC* | Regulator of capsular synthesis sensor histidine kinase | Stimulated by outer membrane damage, LPS synthesis defects, peptidoglycan perturbation, or lipoprotein mislocalisation, it autophosphorilates and transduces the signal to activate the system | 608 | Q52*stop** | [2-5] |
| *aroK* | Shikimate kinase | Aromatic amino acid biosynthesis, possible transcriptional regulator functions | 301 | C36R*<br>E95*stop*<br>59*frameshift* → 91*stop*<br>27*frameshift* → 164*stop* | [6] |
| *tmk* | Thymidylate kinase | Pyrimidine synthesis for DNA synthesis | 208 | E100K* | [7] |
| *ompP* | Outer membrane porin BP0840 | Anion selective channel | 392 | T87S* | [8,9] |
| *lysN* | 2-aminoadipate aminotransferase | Lysine biosynthesis and degradation | 505 | Q466R* | [10] |
| *btuB* | TonB-dependent cobalamin receptor | Outer membrane vitamin B12 receptor | 645 | Full deletion | [11] |
| *btuD* | Cobalamin import ATP-binding protein | ATP-binding cassette component of the vitamin B12 ABC transport system | 500 | Full deletion | [12] |
| *hipA* | Toxin HipA | Serine/threonine kinase toxin of the high incidence of persistence (Hip)AB toxin/antitoxin system | 446 | P254H | [13] |
| *wzb* | Low molecular weight protein-tyrosine-phosphatase | Regulates polysaccharide (glycoconjugate, capsule, Bordetella-specific polysaccharide, exopolysaccharide) chain length by counteracting *wzc* | 163 | M1L (loss of start codon) | [14] |
| *ppgK* | Polyphosphate glucokinase | Phosphorylates glucose into glucose-6-phosphate, which can then be used for many purposes, including LPS and capsule synthesis | 397 | Full deletion | [15] |
| *wecE* | dTDP-4-amino-4,6-dideoxygalactose transaminase | LPS O-antigen nucleotide sugar biosynthesis | 390 | Reversion of 258*frameshift* → 293*stop* to fully functional protein | [16,17] |
| *waaQ* | LPS heptosyltransferase; Glycosyltransferase family 9 protein | LPS core biosynthesis | 325 | T37K<br>L138R<br>S249Y<br>R291P | [18,19] |
| *pgaB* | Polysaccharide deacetylase | Putative role in LPS core synthesis; flanked by two glycosyltransferases | 287 | H93R | [20] |
| *lptD* | LPS assembly protein | LPS assembly at the surface of the outer membrane | 819 | Full deletion | [21] |
| *lptG* | LPS export ABC transporter permease | LPS export to the outer membrane | 392 | K168T | [21] |
| *anmK* | Anhydro-N-acetylmuramic acid kinase | Cell wall biogenesis; peptidoglycan recycling | 372 | 43*frameshift*<br>L312R | [22] |
| *mrcA* | Penicillin-binding protein 1A | DD-transpeptidase, cell wall biogenesis | 868 | Full deletion | [23] |

| | | | | | |
|---|---|---|---|---|---|
| *gmhB* | D-glycero-D-manno-heptose-1,7-bisphosphate phosphatase | Lipid A biosynthesis | 181 | Full deletion | [24] |
| *amgK* | N-acetylmuramate 1-kinase | Peptidoglycan recycling | 356 | Full deletion | [25] |
| *murB* | UDP-N-acetylmuramate dehydrogenase | Peptidoglycan biosynthesis and degradation | 349 | Full deletion | [25] |
| *murU* | N-acetyl-alpha-D-muramate 1-phosphate uridylyltransferase | Peptidoglycan recycling | 230 | Full deletion | [25] |

**Supplementary Note 1.**

**Methodology for antimicrobial susceptibility testing.** Contemporary isolates predominantly underwent initial MIC determination using gradient strip diffusion (bioMérieux; Liofilchem). A non-standard methodology was required given the slow growth of the organism and poor growth on Mueller-Hinton agar; this involved the use of Mueller-Hinton Fastidious agar (Thermofisher, Australia) incubated at 35°C in ambient air and plates read at 48 h of incubation. These results were later confirmed with broth microdilution (Thermofisher, Australia) using standard methodology. Cefiderocol was tested by disc diffusion, fosfomycin was tested by agar dilution, and colistin by broth microdilution.

**Supplementary Note 2.**

**Genomic analysis of phage øSimón.** The complete assembled genome of øSimón was 60,921 bp long, with 62.6% GC content. øSimón had 80 open reading frames (ORFs), no significant matches to previously published phages, and could not be placed into any known genus or family. The closest related phage family was *Mesyanzhinovviridae* (bit-score = 58), known to contain siphoviruses that infect *Bordetella bronchiseptica*[26].

**Supplementary Note 3.**

**Methodology for anti-øSimón depletion antibodies from serum.** Individual wells of MICROLON™ high binding 96-well plates (Greiner, 655061) were coated with 100 μl of a 1:4 solution of phage øSimón in coating buffer (carbonate-bicarbonate with pH = 9.6, Sigma C3041), for approximately at $10^6$ pfu/well. Plates were covered with foil and incubated at 4 °C for 20 h. The wells were emptied, washed 4 times with 300 μl of PBS + 0.05% Tween 80, blocked with 200 μl of 3% skim milk powder in PBS, and the plates incubated as above. The blocking solution was shaken off, and the plates patted dry with paper towels. Next, a 3 ml solution of 1:100 serum in PBS was prepared. An aliquot of 800 μl was saved as the "pre-depletion" sample. The serum solution was split into 100 μl aliquots, each transferred into one of the prepared øSimón-covered wells and left to incubate for 1 h, before transferring it into a fresh øSimón-covered well. After each passage, an aliquot was taken to quantify the anti-øSimón antibody signal through the ELISA methodology described in the main manuscript, corroborating the depletion of the specific antibodies. This was done for 6 passages in a pilot run, and 12 passages in the final runs, with the resulting solution being labelled "post-depletion" sample and used alongside the "pre-depletion" sample for a neutralisation assay as described in the main manuscript. Before the neutralisation assay, the amount of residual øSimón in the post-depletion sample was quantified and later used to correct the calculations.
