## Supplementary Table 4 for "Mechanistic insights and clinical implications of cross-reactive anti-prophage antibodies and bacterial heteroresistance on phage therapeutic failure"

| accession | study | bioproject_accession | biosample_accession | platform | geographic_location | year | species | source | treatment_time_point_(days) |
|---|---|---|---|---|---|---|---|---|---|
| SRR31402293 | SRP542529 | PRJNA1180749 | SAMN44527766 | ILLUMINA | Melbourne, Australia | 2022 | Bordetella bronchialis | Sputum | Day -1 |
| SRR31402292 | SRP542529 | PRJNA1180749 | SAMN44527766 | ILLUMINA | Melbourne, Australia | 2022 | Bordetella bronchialis | Sputum | Day -1 |
| SRR31402281 | SRP542529 | PRJNA1180749 | SAMN44527766 | ILLUMINA | Melbourne, Australia | 2022 | Bordetella bronchialis | Sputum | Day -1 |
| SRR31402259 | SRP542529 | PRJNA1180749 | SAMN44527766 | ILLUMINA | Melbourne, Australia | 2022 | Bordetella bronchialis | Sputum | Day -1 |
| SRR31402248 | SRP542529 | PRJNA1180749 | SAMN44527766 | ILLUMINA | Melbourne, Australia | 2022 | Bordetella bronchialis | Sputum | Day -1 |
| SRR31402237 | SRP542529 | PRJNA1180749 | SAMN44527766 | ILLUMINA | Melbourne, Australia | 2022 | Bordetella bronchialis | Sputum | Day -1 |
| SRR31402270 | SRP542529 | PRJNA1180749 | SAMN44527766 | ILLUMINA | Melbourne, Australia | 2022 | Bordetella bronchialis | Sputum | Day -1 |
| SRR31402269 | SRP542529 | PRJNA1180749 | SAMN44527766 | ILLUMINA | Melbourne, Australia | 2022 | Bordetella bronchialis | Sputum | Day -1 |
| SRR31402268 | SRP542529 | PRJNA1180749 | SAMN44527766 | ILLUMINA | Melbourne, Australia | 2022 | Bordetella bronchialis | Sputum | Day -1 |
| SRR31402267 | SRP542529 | PRJNA1180749 | SAMN44527766 | ILLUMINA | Melbourne, Australia | 2022 | Bordetella bronchialis | Sputum | Day -1 |
| SRR31402291 | SRP542529 | PRJNA1180749 | SAMN44527766 | ILLUMINA | Melbourne, Australia | 2022 | Bordetella bronchialis | Sputum | Day -1 |
| SRR31402290 | SRP542529 | PRJNA1180749 | SAMN44527766 | ILLUMINA | Melbourne, Australia | 2022 | Bordetella bronchialis | Sputum | Day -1 |
| SRR31402289 | SRP542529 | PRJNA1180749 | SAMN44527766 | ILLUMINA | Melbourne, Australia | 2022 | Bordetella bronchialis | Sputum | Day -1 |
| SRR31402288 | SRP542529 | PRJNA1180749 | SAMN44527766 | ILLUMINA | Melbourne, Australia | 2022 | Bordetella bronchialis | Sputum | Day -1 |
| SRR31402287 | SRP542529 | PRJNA1180749 | SAMN44527766 | ILLUMINA | Melbourne, Australia | 2022 | Bordetella bronchialis | Sputum | Day 2 |
| SRR31402286 | SRP542529 | PRJNA1180749 | SAMN44527766 | ILLUMINA | Melbourne, Australia | 2022 | Bordetella bronchialis | Sputum | Day 2 |
| SRR31402285 | SRP542529 | PRJNA1180749 | SAMN44527766 | ILLUMINA | Melbourne, Australia | 2022 | Bordetella bronchialis | Sputum | Day 2 |
| SRR31402284 | SRP542529 | PRJNA1180749 | SAMN44527766 | ILLUMINA | Melbourne, Australia | 2022 | Bordetella bronchialis | Sputum | Day 2 |
| SRR31402283 | SRP542529 | PRJNA1180749 | SAMN44527766 | ILLUMINA | Melbourne, Australia | 2022 | Bordetella bronchialis | Sputum | Day 2 |
| SRR31402282 | SRP542529 | PRJNA1180749 | SAMN44527766 | ILLUMINA | Melbourne, Australia | 2022 | Bordetella bronchialis | Sputum | Day 2 |
| SRR31402280 | SRP542529 | PRJNA1180749 | SAMN44527766 | ILLUMINA | Melbourne, Australia | 2022 | Bordetella bronchialis | Sputum | Day 2 |
| SRR31402279 | SRP542529 | PRJNA1180749 | SAMN44527766 | ILLUMINA | Melbourne, Australia | 2022 | Bordetella bronchialis | Sputum | Day 2 |
| SRR31402278 | SRP542529 | PRJNA1180749 | SAMN44527766 | ILLUMINA | Melbourne, Australia | 2022 | Bordetella bronchialis | Sputum | Day 6 |
| SRR31402266 | SRP542529 | PRJNA1180749 | SAMN44527766 | ILLUMINA | Melbourne, Australia | 2022 | Bordetella bronchialis | Sputum | Day 6 |
| SRR31402265 | SRP542529 | PRJNA1180749 | SAMN44527766 | ILLUMINA | Melbourne, Australia | 2022 | Bordetella bronchialis | Sputum | Day 6 |
| SRR31402264 | SRP542529 | PRJNA1180749 | SAMN44527766 | ILLUMINA | Melbourne, Australia | 2022 | Bordetella bronchialis | Sputum | Day 6 |
| SRR31402263 | SRP542529 | PRJNA1180749 | SAMN44527766 | ILLUMINA | Melbourne, Australia | 2022 | Bordetella bronchialis | Sputum | Day 6 |
| SRR31402262 | SRP542529 | PRJNA1180749 | SAMN44527766 | ILLUMINA | Melbourne, Australia | 2022 | Bordetella bronchialis | Sputum | Day 6 |
| SRR31402261 | SRP542529 | PRJNA1180749 | SAMN44527766 | ILLUMINA | Melbourne, Australia | 2022 | Bordetella bronchialis | Sputum | Day 6 |
| SRR31402260 | SRP542529 | PRJNA1180749 | SAMN44527766 | ILLUMINA | Melbourne, Australia | 2022 | Bordetella bronchialis | Sputum | Day 8 |
| SRR31402258 | SRP542529 | PRJNA1180749 | SAMN44527766 | ILLUMINA | Melbourne, Australia | 2022 | Bordetella bronchialis | Sputum | Day 8 |
| SRR31402257 | SRP542529 | PRJNA1180749 | SAMN44527766 | ILLUMINA | Melbourne, Australia | 2022 | Bordetella bronchialis | Sputum | Day 8 |
| SRR31402256 | SRP542529 | PRJNA1180749 | SAMN44527766 | ILLUMINA | Melbourne, Australia | 2022 | Bordetella bronchialis | Sputum | Day 8 |
| SRR31402255 | SRP542529 | PRJNA1180749 | SAMN44527766 | ILLUMINA | Melbourne, Australia | 2022 | Bordetella bronchialis | Sputum | Day 8 |
| SRR31402254 | SRP542529 | PRJNA1180749 | SAMN44527766 | ILLUMINA | Melbourne, Australia | 2022 | Bordetella bronchialis | Sputum | Day 8 |
| SRR31402253 | SRP542529 | PRJNA1180749 | SAMN44527766 | ILLUMINA | Melbourne, Australia | 2022 | Bordetella bronchialis | Sputum | Day 8 |
| SRR31402252 | SRP542529 | PRJNA1180749 | SAMN44527766 | ILLUMINA | Melbourne, Australia | 2022 | Bordetella bronchialis | Sputum | Day 8 |
| SRR31402251 | SRP542529 | PRJNA1180749 | SAMN44527766 | ILLUMINA | Melbourne, Australia | 2022 | Bordetella bronchialis | Sputum | Day 17 |
| SRR31402250 | SRP542529 | PRJNA1180749 | SAMN44527766 | ILLUMINA | Melbourne, Australia | 2022 | Bordetella bronchialis | Sputum | Day 17 |
| SRR31402249 | SRP542529 | PRJNA1180749 | SAMN44527766 | ILLUMINA | Melbourne, Australia | 2022 | Bordetella bronchialis | Sputum | Day 17 |
| SRR31402247 | SRP542529 | PRJNA1180749 | SAMN44527766 | ILLUMINA | Melbourne, Australia | 2022 | Bordetella bronchialis | Sputum | Day 17 |
| SRR31402246 | SRP542529 | PRJNA1180749 | SAMN44527766 | ILLUMINA | Melbourne, Australia | 2022 | Bordetella bronchialis | Sputum | Day 17 |
| SRR31402245 | SRP542529 | PRJNA1180749 | SAMN44527766 | ILLUMINA | Melbourne, Australia | 2022 | Bordetella bronchialis | Sputum | Day 17 |
| SRR31402244 | SRP542529 | PRJNA1180749 | SAMN44527766 | ILLUMINA | Melbourne, Australia | 2022 | Bordetella bronchialis | Sputum | Day 17 |
| SRR31402243 | SRP542529 | PRJNA1180749 | SAMN44527766 | ILLUMINA | Melbourne, Australia | 2022 | Bordetella bronchialis | Sputum | Day 17 |
| SRR31402242 | SRP542529 | PRJNA1180749 | SAMN44527766 | ILLUMINA | Melbourne, Australia | 2022 | Bordetella bronchialis | Sputum | Day 72 |
| SRR31402241 | SRP542529 | PRJNA1180749 | SAMN44527766 | ILLUMINA | Melbourne, Australia | 2022 | Bordetella bronchialis | Sputum | Day 72 |
| SRR31402240 | SRP542529 | PRJNA1180749 | SAMN44527766 | ILLUMINA | Melbourne, Australia | 2022 | Bordetella bronchialis | Sputum | Day 72 |
| SRR31402239 | SRP542529 | PRJNA1180749 | SAMN44527766 | ILLUMINA | Melbourne, Australia | 2022 | Bordetella bronchialis | Sputum | Day 72 |
| SRR31402238 | SRP542529 | PRJNA1180749 | SAMN44527766 | ILLUMINA | Melbourne, Australia | 2022 | Bordetella bronchialis | Sputum | Day 72 |
| SRR31402236 | SRP542529 | PRJNA1180749 | SAMN44527766 | ILLUMINA | Melbourne, Australia | 2022 | Bordetella bronchialis | Sputum | Day 72 |
| SRR31402235 | SRP542529 | PRJNA1180749 | SAMN44527766 | ILLUMINA | Melbourne, Australia | 2022 | Bordetella bronchialis | Sputum | Day 72 |

| | | | | | | | | | |
|---|---|---|---|---|---|---|---|---|---|
| SRR31402277 | SRP542529 | PRJNA1180749 | SAMN44527766 | ILLUMINA | Melbourne, Australia | 2022 | Bordetella bronchialis | Sputum | Day 84 |
| SRR31402276 | SRP542529 | PRJNA1180749 | SAMN44527766 | ILLUMINA | Melbourne, Australia | 2022 | Bordetella bronchialis | Sputum | Day 84 |
| SRR31402275 | SRP542529 | PRJNA1180749 | SAMN44527766 | ILLUMINA | Melbourne, Australia | 2022 | Bordetella bronchialis | Sputum | Day 84 |
| SRR31402274 | SRP542529 | PRJNA1180749 | SAMN44527766 | ILLUMINA | Melbourne, Australia | 2022 | Bordetella bronchialis | Sputum | Day 84 |
| SRR31402273 | SRP542529 | PRJNA1180749 | SAMN44527766 | ILLUMINA | Melbourne, Australia | 2022 | Bordetella bronchialis | Sputum | Day 84 |
| SRR31402272 | SRP542529 | PRJNA1180749 | SAMN44527766 | ILLUMINA | Melbourne, Australia | 2022 | Bordetella bronchialis | Sputum | Day 84 |
| SRR31189424 | SRP542529 | PRJNA1180749 | SAMN44527766 | ILLUMINA | Melbourne, Australia | 2022 | Bordetella bronchialis | Sputum | NA |
| SRR31189423 | SRP542529 | PRJNA1180749 | SAMN44527766 | OXFORD NANOPORE | Melbourne, Australia | 2022 | Bordetella bronchialis | Sputum | NA |
| CP178749 | NA | PRJNA1180749 | SAMN44527766 | OXFORD NANOPORE, ILLUMINA | Melbourne, Australia | 2022 | Bordetella bronchialis | Sputum | NA |
| SRR31402271 | SRP542529 | PRJNA1180749 | SAMN44527766 | ILLUMINA | Melbourne, Australia | 2022 | Bordetella bronchialis | In vitro obtained, originally sputum | NA |
| PQ683277 | NA | NA | NA | OXFORD NANOPORE, ILLUMINA | Melbourne, Australia | 2022 | Phage Simón | Raw sewage | NA |

| phage_profile | Isolate | format | assembly_method | description |
|---|---|---|---|---|
| No phage selection | 1 | fastq | NA | Illumina WGS of Bordetella bronchialis isolated from CF patient sputum before phage Simon treatment; day -1 isolate 1 |
| No phage selection | 2 | fastq | NA | Illumina WGS of Bordetella bronchialis isolated from CF patient sputum before phage Simon treatment; day -1 isolate 2 |
| No phage selection | 3 | fastq | NA | Illumina WGS of Bordetella bronchialis isolated from CF patient sputum before phage Simon treatment; day -1 isolate 3 |
| No phage selection | 4 | fastq | NA | Illumina WGS of Bordetella bronchialis isolated from CF patient sputum before phage Simon treatment; day -1 isolate 4 |
| No phage selection | 5 | fastq | NA | Illumina WGS of Bordetella bronchialis isolated from CF patient sputum before phage Simon treatment; day -1 isolate 5 |
| No phage selection | 6 | fastq | NA | Illumina WGS of Bordetella bronchialis isolated from CF patient sputum before phage Simon treatment; day -1 isolate 6 |
| No phage selection | 7 | fastq | NA | Illumina WGS of Bordetella bronchialis isolated from CF patient sputum before phage Simon treatment; day -1 isolate 7 |
| No phage selection | 8 | fastq | NA | Illumina WGS of Bordetella bronchialis isolated from CF patient sputum before phage Simon treatment; day -1 isolate 8 |
| No phage selection | 9 | fastq | NA | Illumina WGS of Bordetella bronchialis isolated from CF patient sputum before phage Simon treatment; day -1 isolate 9 |
| No phage selection | 10 | fastq | NA | Illumina WGS of Bordetella bronchialis isolated from CF patient sputum before phage Simon treatment; day -1 isolate 10 |
| No phage selection | 11 | fastq | NA | Illumina WGS of Bordetella bronchialis isolated from CF patient sputum before phage Simon treatment; day -1 isolate 11 |
| No phage selection | 12 | fastq | NA | Illumina WGS of Bordetella bronchialis isolated from CF patient sputum before phage Simon treatment; day -1 isolate 12 |
| No phage selection | 13 | fastq | NA | Illumina WGS of Bordetella bronchialis isolated from CF patient sputum before phage Simon treatment; day -1 isolate 13 |
| No phage selection | 14 | fastq | NA | Illumina WGS of Bordetella bronchialis isolated from CF patient sputum before phage Simon treatment; day -1 isolate 14 |
| Phage-tolerant | 1 | fastq | NA | Illumina WGS of Bordetella bronchialis isolated from CF patient sputum after commencement of phage Simon treatment; day 2 isolate 1 |
| Phage-tolerant | 2 | fastq | NA | Illumina WGS of Bordetella bronchialis isolated from CF patient sputum after commencement of phage Simon treatment; day 2 isolate 2 |
| Phage-tolerant | 3 | fastq | NA | Illumina WGS of Bordetella bronchialis isolated from CF patient sputum after commencement of phage Simon treatment; day 2 isolate 3 |
| Phage-tolerant | 4 | fastq | NA | Illumina WGS of Bordetella bronchialis isolated from CF patient sputum after commencement of phage Simon treatment; day 2 isolate 4 |
| Phage-tolerant | 5 | fastq | NA | Illumina WGS of Bordetella bronchialis isolated from CF patient sputum after commencement of phage Simon treatment; day 2 isolate 5 |
| Phage-tolerant | 6 | fastq | NA | Illumina WGS of Bordetella bronchialis isolated from CF patient sputum after commencement of phage Simon treatment; day 2 isolate 6 |
| Phage-tolerant | 7 | fastq | NA | Illumina WGS of Bordetella bronchialis isolated from CF patient sputum after commencement of phage Simon treatment; day 2 isolate 7 |
| Phage-tolerant | 8 | fastq | NA | Illumina WGS of Bordetella bronchialis isolated from CF patient sputum after commencement of phage Simon treatment; day 2 isolate 8 |
| Phage-tolerant | 1 | fastq | NA | Illumina WGS of Bordetella bronchialis isolated from CF patient sputum after commencement of phage Simon treatment; day 6 isolate 1 |
| Phage-tolerant | 2 | fastq | NA | Illumina WGS of Bordetella bronchialis isolated from CF patient sputum after commencement of phage Simon treatment; day 6 isolate 2 |
| Phage-tolerant | 3 | fastq | NA | Illumina WGS of Bordetella bronchialis isolated from CF patient sputum after commencement of phage Simon treatment; day 6 isolate 3 |
| Phage-tolerant | 4 | fastq | NA | Illumina WGS of Bordetella bronchialis isolated from CF patient sputum after commencement of phage Simon treatment; day 6 isolate 4 |
| Phage-tolerant | 5 | fastq | NA | Illumina WGS of Bordetella bronchialis isolated from CF patient sputum after commencement of phage Simon treatment; day 6 isolate 5 |
| Phage-tolerant | 7 | fastq | NA | Illumina WGS of Bordetella bronchialis isolated from CF patient sputum after commencement of phage Simon treatment; day 6 isolate 7 |
| Phage-tolerant | 8 | fastq | NA | Illumina WGS of Bordetella bronchialis isolated from CF patient sputum after commencement of phage Simon treatment; day 6 isolate 8 |
| Phage-tolerant | 1 | fastq | NA | Illumina WGS of Bordetella bronchialis isolated from CF patient sputum after commencement of phage Simon treatment; day 8 isolate 1 |
| Phage-tolerant | 2 | fastq | NA | Illumina WGS of Bordetella bronchialis isolated from CF patient sputum after commencement of phage Simon treatment; day 8 isolate 2 |
| Phage-tolerant | 3 | fastq | NA | Illumina WGS of Bordetella bronchialis isolated from CF patient sputum after commencement of phage Simon treatment; day 8 isolate 3 |
| Phage-tolerant | 4 | fastq | NA | Illumina WGS of Bordetella bronchialis isolated from CF patient sputum after commencement of phage Simon treatment; day 8 isolate 4 |
| Phage-tolerant | 5 | fastq | NA | Illumina WGS of Bordetella bronchialis isolated from CF patient sputum after commencement of phage Simon treatment; day 8 isolate 5 |
| Phage-tolerant | 6 | fastq | NA | Illumina WGS of Bordetella bronchialis isolated from CF patient sputum after commencement of phage Simon treatment; day 8 isolate 6 |
| Phage-tolerant | 7 | fastq | NA | Illumina WGS of Bordetella bronchialis isolated from CF patient sputum after commencement of phage Simon treatment; day 8 isolate 7 |
| Phage-tolerant | 8 | fastq | NA | Illumina WGS of Bordetella bronchialis isolated from CF patient sputum after commencement of phage Simon treatment; day 8 isolate 8 |
| Phage-tolerant | 1 | fastq | NA | Illumina WGS of Bordetella bronchialis isolated from CF patient sputum after commencement of phage Simon treatment; day 17 isolate 1 |
| Phage-tolerant | 2 | fastq | NA | Illumina WGS of Bordetella bronchialis isolated from CF patient sputum after commencement of phage Simon treatment; day 17 isolate 2 |
| Phage-tolerant | 3 | fastq | NA | Illumina WGS of Bordetella bronchialis isolated from CF patient sputum after commencement of phage Simon treatment; day 17 isolate 3 |
| Phage-tolerant | 4 | fastq | NA | Illumina WGS of Bordetella bronchialis isolated from CF patient sputum after commencement of phage Simon treatment; day 17 isolate 4 |
| Phage-tolerant | 5 | fastq | NA | Illumina WGS of Bordetella bronchialis isolated from CF patient sputum after commencement of phage Simon treatment; day 17 isolate 5 |
| Phage-tolerant | 6 | fastq | NA | Illumina WGS of Bordetella bronchialis isolated from CF patient sputum after commencement of phage Simon treatment; day 17 isolate 6 |
| Phage-tolerant | 7 | fastq | NA | Illumina WGS of Bordetella bronchialis isolated from CF patient sputum after commencement of phage Simon treatment; day 17 isolate 7 |
| Phage-tolerant | 8 | fastq | NA | Illumina WGS of Bordetella bronchialis isolated from CF patient sputum after commencement of phage Simon treatment; day 17 isolate 8 |
| Phage-tolerant | 1 | fastq | NA | Illumina WGS of Bordetella bronchialis isolated from CF patient sputum after commencement of phage Simon treatment; day 72 isolate 1 |
| Phage-tolerant | 2 | fastq | NA | Illumina WGS of Bordetella bronchialis isolated from CF patient sputum after commencement of phage Simon treatment; day 72 isolate 2 |
| Phage-tolerant | 3 | fastq | NA | Illumina WGS of Bordetella bronchialis isolated from CF patient sputum after commencement of phage Simon treatment; day 72 isolate 3 |
| Phage-tolerant | 4 | fastq | NA | Illumina WGS of Bordetella bronchialis isolated from CF patient sputum after commencement of phage Simon treatment; day 72 isolate 4 |
| Phage-tolerant | 5 | fastq | NA | Illumina WGS of Bordetella bronchialis isolated from CF patient sputum after commencement of phage Simon treatment; day 72 isolate 5 |
| Phage-tolerant | 6 | fastq | NA | Illumina WGS of Bordetella bronchialis isolated from CF patient sputum after commencement of phage Simon treatment; day 72 isolate 6 |
| Phage-tolerant | 7 | fastq | NA | Illumina WGS of Bordetella bronchialis isolated from CF patient sputum after commencement of phage Simon treatment; day 72 isolate 7 |

| | | | | |
|---|---|---|---|---|
| Phage-tolerant | 1 | fastq | NA | Illumina WGS of Bordetella bronchialis isolated from CF patient sputum after commencement of phage Simon treatment; day 84 isolate 1 |
| Phage-tolerant | 3 | fastq | NA | Illumina WGS of Bordetella bronchialis isolated from CF patient sputum after commencement of phage Simon treatment; day 84 isolate 3 |
| Phage-tolerant | 4 | fastq | NA | Illumina WGS of Bordetella bronchialis isolated from CF patient sputum after commencement of phage Simon treatment; day 84 isolate 4 |
| Phage-tolerant | 5 | fastq | NA | Illumina WGS of Bordetella bronchialis isolated from CF patient sputum after commencement of phage Simon treatment; day 84 isolate 5 |
| Phage-tolerant | 6 | fastq | NA | Illumina WGS of Bordetella bronchialis isolated from CF patient sputum after commencement of phage Simon treatment; day 84 isolate 6 |
| Phage-tolerant | 7 | fastq | NA | Illumina WGS of Bordetella bronchialis isolated from CF patient sputum after commencement of phage Simon treatment; day 84 isolate 7 |
| Phage sensitive | NA | fastq | NA | Illumina WGS of WT Bordetella bronchialis isolated from CF patient sputum |
| Phage sensitive | NA | fastq | NA | ONT WGS of WT Bordetella bronchialis isolated from CF patient sputum |
| Phage sensitive | NA | fasta | Trycycler v0.5.4 | Complete genomic assembly of WT Bordetella bronchialis isolated from CF patient sputum |
| Phage resistant | NA | fastq | NA | Illumina WGS of phage Simón-resistant Bordetella bronchialis isolated from lab |
| NA | NA | fasta | Unicycler | Genomic assembly of Phage Simon isolated from raw sewage |
