## Supplementary File 1 for "Mechanistic insights and clinical implications of cross-reactive anti-prophage antibodies and bacterial heteroresistance on phage therapeutic failure"

### Day -1

| Position | Gene Code | Gene | Consequence | Reference | 1 | 2 | 3 | 4 | 5 | 6 | 7 | 8 | 9 | 10 | 11 | 12 | 13 | 14 | Strain |
|---|---|---|---|---|---|---|---|---|---|---|---|---|---|---|---|---|---|---|---|
| | | | | | 1 | 2 | 2 | 1 | 1 | 2 | 2 | 2 | 2 | 2 | 2 | 2 | 2 | 2 | Cluster |
| | | | | | 1.68 | 1.64 | 1.231 | 1.205 | 1.042 | 0.913 | 0.908 | 0.841 | 0.758 | 0.712 | 0.692 | 0.665 | 0.537 | 0.484 | Median EOP |
| 155369 | | Intergenic | | C | | | | | | | A | | | | | | | | |
| 167924 | CIIBHN_00745 | Glutathione S-transferase | P81L | G | | | | | | | | | | | | | | A | |
| 202413 | CIIBHN_00925 | glutamate synthase large subunit | T21M | G | | | | | | | | | | | | | A | | |
| 210734 | CIIBHN_00960 | Sodium:proton exchanger | Y668* | C | | | | | | | | | | | | | | G | |
| 236439 | | Intergenic | | T | | C | C | | | | | | | | | | | | |
| 251903 | CIIBHN_01175 | 3-dmu-9-3-mt domain-containing protein | P66R | C | | | | | G | | | | | | | | | | |
| 338354 | CIIBHN_01505 | 5-oxoprolinase | E427A | A | | | | | | | | | | | | | | C | |
| 350851 | | Intergenic | | C | | | | | | | | | T | | | | | | |
| 360689 | CIIBHN_01620 | Alcohol dehydrogenase | Q49P | A | | | | | | | | | | | | | | C | |
| 363559 | | Intergenic | | T | | A | A | | | A | A | A | A | A | A | A | A | A | |
| 529484 | CIIBHN_02365 | RNA methyltransferase | Synonymous | C | | | | | | | G | | | | | | | | |
| 542046 | CIIBHN_02435 | META domain-containing protein | Synonymous | T | | C | C | | | C | C | C | C | C | C | C | C | C | |
| 553457 | CIIBHN_02475 | DUF490 domain-containing protein | V1001G | T | G | | | G | | | | | | | | | | | |
| 579697 | | Intergenic | | T | C | | | C | | | C | C | C | C | | C | C | C | |
| 579703 | | Intergenic | | C | | | | | | | | | | | | | | A | |
| 609961 | | Intergenic | | G | | A | A | | | | | | | | | | | | |
| 658050 | CIIBHN_02920 | Malate dehydrogenase | Synonymous | C | | | | | | T | | | | | | | | | |
| 683477..<br>683944 | CIIBHN_03020 | HTH tetR-type domain-containing protein<br>(transcriptional regulator) | * | | | | | IS481 | IS481 | | | IS481 | | IS481 | | IS481 | IS481 | IS481 | |
| 684990..<br>686186 | CIIBHN_03030 | LysR substrate-binding domain-containing protein<br>(transcriptional regulator) | * | | | | | IS481 | IS481 | | | IS481 | | IS481 | | IS481 | IS481 | IS481 | |
| 933348 | CIIBHN_04205 | Transcriptional regulator, possible shikimate kinase | C36R | T | | C | C | | | C | C | C | C | C | C | C | C | C | |
| 933525 | CIIBHN_04205 | Transcriptional regulator, possible shikimate kinase | E95* | G | | | | | | T | | | | | | | | | |
| 961292 | CIIBHN_04315 | formyltetrahydrofolate deformylase | A84V | G | | | | | | | | | | | | | A | | |
| 1052074 | | Intergenic | | T | | | | | | | | | | | | C | | | |
| 1092510 | | Intergenic | | T | | | | | | A | A | A | A | A | A | A | A | | |
| 1220753 | CIIBHN_05525 | dihydropteroate synthase | P66S | C | T | | | T | | | | | | | | | | | |
| 1308456 | | Intergenic | | G | | | | | C | | | | | | | | | | |
| 1371605 | CIIBHN_06185 | Carnitine dehydratase | A340P | C | | | | | | | | | | | | | | G | |
| 1401888 | | Intergenic | | C | | | G | | | | | | | | | | | | |
| 1449143 | CIIBHN_06550 | Ligand-gated channel protein | Synonymous | C | | | | | T | | | | | | | | | | |
| 1542036 | CIIBHN_06950 | aconitate hydratase AcnA | Synonymous | G | | | | | | | | | | | | | C | | |
| 1549571..<br>1550377 | CIIBHN_06985 | RsxB: Electron transport complex subunit | * | | | | | | | | | | Deleted | | | | | | |
| 1565907 | CIIBHN_07065 | Transcriptional regulator | Synonymous | C | | | | | | | | | T | | | | | | |
| 1703774 | CIIBHN_07715 | endopeptidase La | K392E | T | | | | | | C | C | C | C | C | C | C | C | | |
| 1704811 | CIIBHN_07715 | endopeptidase La | L46P | A | | | | | | | | | | | | | | G | |
| 1738365 | CIIBHN_07860 | RHS repeat-associated core domain-containing protein | W167* | C | | | | | | T | | | | | | | | | |
| 1775164 | CIIBHN_08000 | alanine--tRNA ligase | Synonymous | C | | | | T | | | | | | | | | | | |
| 1802586 | CIIBHN_08120 | RNA polymerase sigma factor RpoD | R123C | C | | | | | | | | | T | | | | | | |
| 1823405 | CIIBHN_08215 | 2-aminoadipate aminotransferase, lysN | Q466R | T | | C | C | | | C | C | C | C | C | C | C | C | C | |
| 1824452 | CIIBHN_08215 | 2-aminoadipate aminotransferase, lysN | D117V | T | | A | A | | | | | | | | | | | | |
| 1873214 | CIIBHN_08475 | YdiU family protein (selO - adenylyl transferase) | W202L | G | | | | | | | | | | T | | | | | |
| 2029057 | | Intergenic | | G | | | | | | | | | | | | | A | | |
| 2034824 | CIIBHN_09230 | MucB-RseB domain-containing protein | I153V | A | G | G | G | G | G | G | G | G | G | G | G | G | G | G | |
| 2157639 | CIIBHN_09715 | TonB-dependent siderophore receptor | L161R | A | C | | | C | | | | | | | | | | | |
| 2263781 | CIIBHN_10130 | MmgE/PrpD family protein | L250F | C | | | | | | | T | T | T | T | T | T | T | | |
| 2381581 | CIIBHN_10670 | outer membrane protein assembly factor BamA | L147F | C | | | | | | | | | | | | | | T | |
| 2382695 | CIIBHN_10670 | outer membrane protein assembly factor BamA | T518R | C | | | | | | G | G | G | G | G | G | G | G | | |
| 2439802 | CIIBHN_10965 | rcsC: regulator of capsular synthesis sensor histidine kinase | Q52* | C | | T | T | | | T | T | T | T | T | T | T | T | T | |
| 2518442 | | Intergenic | | A | | | | | | | | | | | | | C | | |
| 2630309 | CIIBHN_11785 | DUF802 domain-containing protein | Synonymous | G | | | | | | | C | C | C | C | C | C | C | | |
| 2882051..<br>2882707 | CIIBHN_12950 | hypothetical protein | * | | | | | | IS256 | | | | | | | | | | |
| 3000537 | CIIBHN_13470 | Awr type III effector family protein | S639N | G | | | A | | | | | | | | | | | | |
| 3015385 | CIIBHN_13530 | Twin-arginine translocation pathway signal protein | P269L | G | | | | | A | | | | | | | | | | |
| 3036263 | CIIBHN_13620 | YicC family protein | Synonymous | C | | | | | T | | | | | | | | | | |
| 3078043 | CIIBHN_13805 | GntR family transcriptional regulator | L16H | T | | | | | | | | | A | | | | | | |
| 3341289 | CIIBHN_14935 | DUF1109 domain-containing protein | Y176* | A | C | | | C | | | | | | | | | | | |
| 3376565 | CIIBHN_15100 | hypothetical protein | G185S | G | | A | A | | | | | | | | | | | | |
| 3419008 | CIIBHN_15280 | wzb: low molecular weight protein-tyrosine-phosphatase | T123A | T | | C | | | | | | | | | | | | | |

Day -1

| | | | | | | | | | | | | | | | | | | |
|---|---|---|---|---|---|---|---|---|---|---|---|---|---|---|---|---|---|---|
| 3419368 | CIIBHN_15280 | wzb: low molecular weight protein-tyrosine-phosphatase | T3A | T | | | C | | | | | | | | | | | |
| 3445552 | CIIBHN_15390 | isoleucine--tRNA ligase | P114A | C | | | | | | | G | | | | | | | |
| 3454974 | CIIBHN_15435 | Nramp family divalent metal transporter | Synonymous | C | | | | | | | | | | | | | | T |
| 3622057 | CIIBHN_16190 | Spermidine/putrescine ABC transporter permease | Synonymous | G | | | | | | | | | | | | | A | |
| 3631374 | CIIBHN_16240 | ABC transporter substrate-binding protein | Synonymous | T | | | | | | | | | | | C | | | |
| 3638231 | CIIBHN_16275 | Acyl-CoA dehydrogenase | Synonymous | G | A | | | | | | | | | | | | | |
| 3837396 | CIIBHN_17390 | Electron transfer flavoprotein subunit beta | P215S | G | | | | | | | | | A | | | | | |
| 3853445 | CIIBHN_17470 | prephenate dehydratase | D279Y | C | | | | | | | | | | | | | A | |
| 3853474 | CIIBHN_17470 | prephenate dehydratase | G269D | C | | | | | | | | | T | | | | | |
| 3854040 | CIIBHN_17470 | prephenate dehydratase | W80C | C | | | | | | | | | | A | | | | |
| 3857801 | CIIBHN_17480 | DNA gyrase subunit A | N87Y | T | | A | A | | | | | | | | | | | |
| 3861399… 3907354 | CIIBHN_17505 CIIBHN_17825 | Prophage | * | | Deleted | | | Deleted | | | | | | | | | | |
| 3926090 | CIIBHN_17915 | 3-deoxy-7-phosphoheptulonate synthase | G231S | G | | | | | | | | | | | | | | A |
| 3937197 | CIIBHN_17970 | Chemotaxis protein cheW (KEGG Ortholog of WspB) | A80V | C | | | | | | | T | T | | | T | T | | |
| 3999174 | CIIBHN_18155 | citrate synthase | E410D | C | G | G | G | G | | G | G | G | G | G | G | G | G | G |
| 4037531 | | Intergenic | | T | | | | | A | | | | | | | | | |
| 4037532 | | Intergenic | | G | | | | | T | | | | | | | | | |
| 4040180.. 4041070 | CIIBHN_18335 | LysR family transcriptional regulator | * | | | | IS481 | | | | | | | | | | | |
| 4041260.. 4044328 | CIIBHN_18340 | formate dehydrogenase-N subunit alpha | * | | | | IS481 | | | | | | | | | | | |
| 4059158 | CIIBHN_18415 | PBPb domain-containing protein | Synonymous | C | | | | | T | | | | | | | | | |
| 4108696 | CIIBHN_18625 | Outer membrane porin protein BP0840, OmpP | T87S | T | | A | A | | | A | A | A | A | A | A | A | A | A |
| 4233634 | CIIBHN_19235 | nucleotidyltransferase | Synonymous | G | T | | | T | | | | | | | | | | |
| 4233678 | CIIBHN_19235 | nucleotidyltransferase | L265F | G | A | | | A | | | | | | | | | | |
| 4328013 | | Intergenic | | A | | G | G | | | G | G | G | G | G | G | G | G | G |
| 4328024 | | Intergenic | | G | | A | A | | | A | A | A | A | A | A | A | A | A |
| 4364236 | CIIBHN_19780 | hypothetical protein | M37V | A | G | G | G | G | G | G | G | G | G | G | G | G | G | G |
| 4368810 | CIIBHN_19795 | Amino acid deaminase | K357E | A | G | G | G | G | | G | G | G | G | G | G | G | G | G |
| 4420806.. 4421057 | CIIBHN_20025 | hypothetical protein (truncated filamentous haemagglutinin N-terminal domain) | * | | | IS256 | | | | | | | | | | | IS256 | |
| 4424326.. 4425633 | CIIBHN_20035 | citrate synthase (unknown stereospecificity) | * | | | IS256 | | | | | | | | | | | IS256 | |
| 4534006 | CIIBHN_20530 | type III secretion system export apparatus subunit SctV (bcrD, yscV, escV) | D598Y | G | T | T | T | T | | T | T | T | T | T | T | T | T | T |
| 4584761 | | Intergenic | | C | | T | T | | | | | | | | | | | |
| 4670782 | CIIBHN_21155 | GntR family transcriptional regulator | Synonymous | G | | A | A | | | | | | | | | | | |
| 4697533 | CIIBHN_21255 | Hydrolase | A21V | C | | T | | | | | | | | | | | | |
| 4851242 | CIIBHN_21875 | Phage-base-V domain-containing protein | I611N | T | | A | A | | | | | | | | | | | |
| 4851259 | CIIBHN_21875 | Phage-base-V domain-containing protein | G617R | G | C | | | C | | | | | | | | | | |
| 4902525 | CIIBHN_22095 | DUF4232 domain-containing protein | G108D | G | A | | | A | | | | | | | | | | |
| 4939176 | CIIBHN_22235 | MFS transporter | C534* | C | A | | | | | | | | | | | | | |
| 4977906.. 4979378 | CIIBHN_22410 | hydA: dihydropyrimidinase | * | | | | | | | | | | | | | | | Deleted |
| 4979756 | CIIBHN_22415 | transporter substrate-binding domain-containing protein | * | | | | | | | | | | | | | | | Deleted |
| 5029472 | | Intergenic | | T | | | | | | | | | C | | | | | |
| 5062846 | CIIBHN_22785 | Cation-binding protein | T38A | A | G | | | | | | | | | | | | | |
| 5483236 | CIIBHN_24665 | ABC transporter | P158L | G | | A | | | | | | | | | | | | |
| 5641551 | CIIBHN_25340 | Branched chain amino acid ABC transporter substrate-binding protein | Synonymous | T | | C | C | | | C | C | C | C | C | C | C | C | C |
| 5681646 | CIIBHN_25545 | Carnitine dehydratase | S98A | T | | G | G | | | | | | | | | | | |
| 5709159 | CIIBHN_25665 | Thymidylate kinase | E100K | G | | A | A | | | A | A | A | A | A | A | A | A | A |
| 5709579 | CIIBHN_25665 | Thymidylate kinase | N240D | A | | G | G | | | G | G | G | G | G | G | G | G | G |
| 5726613 | | Intergenic | | A | C | | | | | | C | | | | | | | |
| 5771137 | CIIBHN_25995 | DNA-directed RNA polymerase subunit beta' | P1346L | G | A | A | A | A | | A | A | A | A | A | A | A | A | A |
| 5774839 | CIIBHN_25995 | DNA-directed RNA polymerase subunit beta' | A112V | G | | A | A | | | | | | | | | | | |

Day 2

| Position | Gene Code | Gene | Consequence | Reference | 1 | 2 | 3 | 4 | 5 | 6 | 7 | 8 | Strain |
|---|---|---|---|---|---|---|---|---|---|---|---|---|---|
| | | | | | 1 | 1 | 2 | 2 | 2 | 2 | 2 | 1 | Cluster |
| | | | | | 1.4632 | 0.7449 | 0.7279 | 0.5034 | 0.432 | 0.425 | 0.384 | 0 | Median EOP |
| 112720 | CIIBHN_00500 | Acetyl-CoA acetyltransferase | G305S | C | | | | T | | | | | |
| 228455.. 229177 | CIIBHN_01060 | Rick-17kDa-Anti domain-containing protein | * | | | | | Deleted | | | | | |
| 363559 | | Intergenic | | T | | | A | A | A | A | A | | |
| 438841 | CIIBHN_01935 | LPS heptosyltransferase; Glycosyltransferase family 9 protein | L138R | T | | | | | | G | | | |
| 542046 | CIIBHN_02435 | META domain-containing protein | Synonymous | T | | | C | C | C | C | C | | |
| 553457 | CIIBHN_02475 | DUF490 domain-containing protein | V1101G | T | G | | | | | | | | |
| 579697 | | Intergenic | | T | C | | C | C | C | C | C | | |
| 579705 | | Intergenic | | GTT | | | | GTTTT | GTTTT | | | | |
| 683477.. 683944 | CIIBHN_03020 | HTH tetR-type domain-containing protein (transcriptional regulator) | * | | | | IS481 | | IS481 | IS481 | IS481 | IS481 | |
| 684990.. 686186 | CIIBHN_03030 | LysR substrate-binding domain-containing protein (transcriptional regulator) | * | | | | IS481 | | IS481 | IS481 | IS481 | IS481 | |
| 706563 | CIIBHN_03120 | 4,5-dioxygenase | Synonymous | G | | | | | | | T | | |
| 933348 | CIIBHN_04205 | Transcriptional regulator, possible shikimate kinase | C36R | T | | | C | C | C | C | C | | |
| 1092510 | | Intergenic | | T | | | A | A | A | A | A | | |
| 1149449 | CIIBHN_05150 | DNA mismatch repair endonuclease MutL | A75E | G | | | | | | T | | | |
| 1220753 | CIIBHN_05525 | dihydropteroate synthase | P66S | C | T | | | | | | | | |
| 1383568 | CIIBHN_06260 | 4-hydroxy-tetrahydrodipicolinate synthase | S112F | C | | | | | | | | T | |
| 1687807.. 1688760 | CIIBHN_07640 | OmpR/PhoB-type domain-containing protein (transcriptional regulator) | * | | | | | | | | Deleted | | |
| 1703774 | CIIBHN_07715 | endopeptidase La | K392E | T | | | C | C | C | C | C | | |
| 1762069 | CIIBHN_07945 | CDP-6-deoxy-delta-3,4-glucoseen reductase | Synonymous | C | | | | | | T | | | |
| 1823405 | CIIBHN_08215 | 2-aminoadipate aminotransferase, lysN | Q466R | T | | | C | C | C | C | C | | |
| 2034824 | CIIBHN_09230 | MucB-RseB domain-containing protein | I153V | A | G | G | G | G | G | G | G | G | |
| 2074231 | CIIBHN_09405 | ATP-dependent DNA helicase | A1008S | C | | | | | | | | A | |
| 2157639 | CIIBHN_09715 | TonB-dependent siderophore receptor | L161R | A | C | | | | | | | | |
| 2217187 | CIIBHN_09965 | wecE: dTDP-4-amino-4,6-dideoxygalactose transaminase 2217081..2217962 | Frameshift AA 259/293 | TGG | | | | | | | TG | | |
| 2263781 | CIIBHN_10130 | MmgE/PrpD family protein | L250F | C | | | T | T | T | T | T | | |
| 2382695 | CIIBHN_10670 | outer membrane protein assembly factor BamA | T518R | C | | | G | G | G | G | G | | |
| 2439802 | CIIBHN_10965 | rcsC: regulator of capsular synthesis sensor histidine kinase | Q52* | C | | | T | T | T | T | T | | |
| 2470157 | CIIBHN_11090 | Efflux transporter periplasmic adaptor subunit | L388Q | A | | T | | | | | | | |
| 2495405 | CIIBHN_11205 | Glyoxalase | W73R | A | | | | | | T | | | |
| 2630309 | CIIBHN_11785 | DUF802 domain-containing protein | Synonymous | G | | | C | C | C | C | C | | |
| 3294158 | CIIBHN_14675 | ABC-type glycerol-3-phosphate transport system, permease component | I21L | T | | | | | | | | G | |
| 3341289 | CIIBHN_14935 | DUF1109 domain-containing protein | Y176W | A | C | | | | | | | | |
| 3341290 | | | | T | C | | | | | | | | |
| 3450003 | CIIBHN_15405 | Hydrolase | Synonymous | G | | | | | | | C | | |
| 3551595 | CIIBHN_15880 | 2-hydroxy-acid oxidase | Synonymous | G | | A | | | | | | | |
| 3600110 | CIIBHN_16100 | Polysaccharide deacetylase | H93R | T | | C | | | | | | | |
| 3761136 | CIIBHN_16895 | Phage head morphogenesis protein | | T | | | | | | C | | | |
| 3773048 | | Intergenic | | G | | | | | | | | T | |
| 3852400 | CIIBHN_17465 | histidinol-phosphate transaminase | V244A | A | | | | | | G | | | |
| 3854018 | CIIBHN_17470 | prephenate dehydratase | R88C | G | | | | | | | A | | |
| 3854040 | CIIBHN_17470 | prephenate dehydratase | W80C | C | | | | A | A | | | | |
| 3861399… 3907354 | CIIBHN_17505 CIIBHN_17825 | Prophage | * | | Deleted | | | | | | | | |
| 3937197 | CIIBHN_17970 | Chemotaxis protein cheW (KEGG Ortholog of WspB) | A80V | C | | | T | | | | | | |
| 3999174 | CIIBHN_18155 | citrate synthase | E410D | C | G | | G | G | G | G | G | | |
| 4108696 | CIIBHN_18625 | Outer membrane porin protein BP0840, OmpP | T87S | T | | | A | A | A | A | A | | |
| 4233634 | CIIBHN_19235 | nucleotidyltransferase | Synonymous | G | T | | | | | | | | |
| 4233678 | CIIBHN_19235 | nucleotidyltransferase | L265F | G | A | | | | | | | | |
| 4328013 | | Intergenic | | A | | | G | G | G | G | G | | |
| 4328024 | | Intergenic | | G | | | A | A | A | A | A | | |
| 4364236 | CIIBHN_19780 | hypothetical protein | M37V | A | G | G | G | G | G | G | G | G | |
| 4368810 | CIIBHN_19795 | Amino acid deaminase | K357E | A | G | | G | G | | G | G | | |
| 4420806.. 4421057 | CIIBHN_20025 | hypothetical protein (truncated filamentous haemagglutinin N-terminal domain) | * | | IS256 | IS256 | | | | | | | |
| 4424326.. 4425633 | CIIBHN_20035 | citrate synthase (unknown stereospecificity) | * | | IS256 | IS256 | | | | | | | |
| 4513998 | | Intergenic | | G | A | | | | | | | | |
| 4534006 | CIIBHN_20530 | type III secretion system export apparatus subunit SctV (bcrD, yscV, escV) | D598Y | G | T | | T | T | T | T | T | | |
| 4833824 | CIIBHN_21825 | BTAD domain-containing protein | G819S | C | | | | | | | | T | |
| 4851259 | CIIBHN_21875 | Phage-base-V domain-containing protein | G617R | G | C | | | | | | | | |

| | | | | | | | | | | | | |
|---|---|---|---|---|---|---|---|---|---|---|---|---|
| 4902525 | CIIBHN_22095 | DUF4232 domain-containing protein | G108D | G | A | | | | | | | |
| 4913078 | CIIBHN_22130 | Type VI secretion-associated protein | P100L | C | | | | | | | | T |
| 5051393 | CIIBHN_22720 | YbhB/YbcL family Raf kinase inhibitor-like protein | Synonymous | C | | G | | | | | | |
| 5129214 | CIIBHN_23090 | Sensor protein QseC | A103T | G | | | | | | A | | |
| 5230900 | | Intergenic | | G | | | | | | | | A |
| 5290080.. 5293862 | CIIBHN_23785 | metH: methionine synthase | * | | | | | | | | | Deleted |
| 5294266.. 5296203 | CIIBHN_23790 | BtuB: tonB-dependent receptor | * | | | | | | | | | Deleted |
| 5514067 | CIIBHN_24785 | ABC transporter permease | T351A | T | | | | | | C | | |
| 5641551 | CIIBHN_25340 | Branched chain amino acid ABC transporter substrate-binding protein | Synonymous | T | | | | C | C | C | C | |
| 5671718 | CIIBHN_25495 | Toxin HipA (type II toxin-antitoxin system) | P254H | C | A | | | | | | | |
| 5709159 | CIIBHN_25665 | Thymidylate kinase | E100K | G | | | A | A | A | A | A | |
| 5709579 | CIIBHN_25665 | Thymidylate kinase | N240D | A | | | G | G | G | G | G | |
| 5771137 | CIIBHN_25995 | DNA-directed RNA polymerase subunit beta' | P1346L | G | A | | A | A | A | A | A | |

| Position | Gene Code | Gene | Consequence | Reference | 1 | 2 | 3 | 4 | 5 | 6 | 7 | Strain |
|---|---|---|---|---|---|---|---|---|---|---|---|---|
| | | | | | 2 | 2 | 2 | 2 | 2 | 2 | 2 | Cluster |
| | | | | | 0.663 | 0.602 | 0.511 | 0.4853 | 0.3118 | 0.2993 | 0.0272 | Median EOP |
| 32825 | CIIBHN_00140 | LysR substrate-binding domain-containing protein | V148L | G | | C | | | | | | |
| 208884 | CIIBHN_00960 | Sodium:proton exchanger 208731..211241 | Frameshift AA 52/836 (84*) | GC | | | | | | | GCC | |
| 223185 | CIIBHN_01020 | cytochrome-c oxidase | G198S | G | | | | | | | A | |
| 236439 | | Intergenic | | T | C | | | | | | | |
| 363559 | | Intergenic | | T | A | A | A | A | A | A | A | |
| 438538 | CIIBHN_01935 | LPS heptosyltransferase; Glycosyltransferase family 9 protein | T37K | C | | | | | | | A | |
| 439300 | CIIBHN_01935 | LPS heptosyltransferase; Glycosyltransferase family 9 protein | R291P | G | | | | | C | | | |
| 449356 | CIIBHN_01975 | Cytochrome ubiquinol oxidase subunit I | R11H | C | | | | | T | | | |
| 491332 | | Intergenic | | G | | | | | A | | | |
| 542046 | CIIBHN_02435 | META domain-containing protein | Synonymous | T | C | C | C | C | C | C | C | |
| 553445 | CIIBHN_02435 | META domain-containing protein | D1097V | A | T | | | | | | | |
| 579697 | | Intergenic | | T | | | C | C | C | C | C | |
| 609961 | | Intergenic | | G | A | | | | | | | |
| 658050 | CIIBHN_02920 | Malate dehydrogenase | Synonymous | C | | T | | | | | | |
| 683477.. 683944 | CIIBHN_03020 | HTH tetR-type domain-containing protein (transcriptional regulator) | * | | | | | | IS481 | IS481 | IS481 | |
| 684990.. 686186 | CIIBHN_03030 | LysR substrate-binding domain-containing protein (transcriptional regulator) | * | | | | | | IS481 | IS481 | IS481 | |
| 706563 | CIIBHN_03120 | 4,5-dioxygenase | Synonymous | G | | | | | | | T | |
| 933348 | CIIBHN_04205 | Transcriptional regulator, possible shikimate kinase | C36R | T | C | C | C | C | C | C | C | |
| 933419 | CIIBHN_04205 | Transcriptional regulator, possible shikimate kinase | Frameshift AA 59/301 (91*) | TG | T | | | | | | | |
| 1092510 | | Intergenic | | T | | A | A | A | A | A | A | |
| 1159279... 1195971 | CIIBHN_05200 CIIBHN_05410 | Prophage | * | | | Deleted | | | | | | |
| 1460678 | CIIBHN_06605 | histidine kinase | Q204L | A | | | | T | | | | |
| 1486040 | | Intergenic | | T | | | | | | | G | |
| 1687807.. 1688760 | CIIBHN_07640 | OmpR/PhoB-type domain-containing protein (transcriptional regulator) | * | | | | Deleted | | | Deleted | | |
| 1703774 | CIIBHN_07715 | endopeptidase La | K392E | T | | C | C | C | C | C | C | |
| 1823405 | CIIBHN_08215 | 2-aminoadipate aminotransferase, lysN | Q466R | T | C | C | C | C | C | C | C | |
| 1824452 | CIIBHN_08215 | 2-aminoadipate aminotransferase, lysN | D117V | T | A | | | | | | | |
| 2020569 | CIIBHN_09170 | Ligand-gated channel protein | A9T | G | | | | | | A | | |
| 2034824 | CIIBHN_09230 | MucB-RseB domain-containing protein | I153V | A | G | G | G | G | G | G | G | |
| 2091669 | CIIBHN_09485 | HTH lysR-type domain-containing protein | Synonymous | C | | | | | T | | | |
| 2217187 | CIIBHN_09965 | wecE: dTDP-4-amino-4,6-dideoxygalactose transaminase 2217081..2217962 | Frameshift AA 259/293 | TGG | | | TG | | | | | |
| 2263781 | CIIBHN_10130 | MmgE/PrpD family protein | L250F | C | | | T | T | T | T | T | |
| 2286579 | CIIBHN_10215 | folD: bifunctional methylenetetrahydrofolate dehydrogenase methenyltetrahydrofolate cyclohydrolase | T184S | T | A | | | | | | | |
| 2301995 | | Intergenic | | C | | | | T | | | | |
| 2382695 | CIIBHN_10670 | outer membrane protein assembly factor BamA | T518R | C | | G | G | G | G | G | G | |
| 2439802 | CIIBHN_10965 | rcsC: regulator of capsular synthesis sensor histidine kinase | Q52* | C | T | T | T | T | T | T | T | |
| 2630309 | CIIBHN_11785 | DUF802 domain-containing protein | Synonymous | G | | | C | C | C | C | C | |
| 2796807 | CIIBHN_12585 | EF-hand domain-containing protein | A53E | C | | | | | | | A | |
| 2830932 | CIIBHN_12725 | citrate synthase (unknown stereospecificity) | F42C | T | | | | | | | G | |
| 2856929 | CIIBHN_12835 | cytochrome c oxidase accessory protein CcoG | L164P | T | | | | | | C | | |
| 3347375 | CIIBHN_14970 | Methyl-accepting chemotaxis sensory transducer | G468D | G | | | | A | | | | |
| 3376565 | CIIBHN_15100 | hypothetical protein | G185S | G | A | | | | | | | |
| 3522600 | | Intergenic | | A | | G | | | | | | |
| 3854018 | CIIBHN_17470 | prephenate dehydratase | R88C | G | | | | | | | A | |
| 3854040 | CIIBHN_17470 | prephenate dehydratase | W80C | C | | | | | A | A | | |
| 3857801 | CIIBHN_17480 | DNA gyrase subunit A | N87Y | T | A | | | | | | | |
| 3937197 | CIIBHN_17970 | Chemotaxis protein cheW (KEGG Ortholog of WspB) | A80V | C | | | T | T | | | | |
| 3999174 | CIIBHN_18155 | citrate synthase | E410D | C | G | G | G | G | G | G | G | |
| 4007084 | CIIBHN_18195 | 2-methylcitrate synthase | R259H | G | | A | | | | | | |
| 4040180.. 4041070 | CIIBHN_18335 | LysR family transcriptional regulator | * | | IS481 | | | | | | | |
| 4041260.. 4044328 | CIIBHN_18340 | formate dehydrogenase-N subunit alpha | * | | IS481 | | | | | | | |
| 4077437.. 4078498 | CIIBHN_18490 | Type-4 uracil-DNA glycosylase | * | | | | | | | | Deleted | |
| 4108696 | CIIBHN_18625 | Outer membrane porin protein BP0840, OmpP | T87S | T | A | A | A | A | A | A | A | |
| 4328013 | | Intergenic | | A | G | G | G | G | G | G | G | |
| 4328024 | | Intergenic | | G | A | A | A | A | A | A | A | |

Day 6

| | | | | | | | | | | | |
|---|---|---|---|---|---|---|---|---|---|---|---|
| 4364236 | CIIBHN_19780 | hypothetical protein | M37V | A | G | G | G | G | G | G | G |
| 4368810 | CIIBHN_19795 | Amino acid deaminase | K357E | A | G | G | G | G | G | G | G |
| 4420806..<br>4421057 | CIIBHN_20025 | hypothetical protein<br>(truncated filamentous haemagglutinin N-terminal domain) | * | | IS256 | IS256 | | | | | |
| 4424326..<br>4425633 | CIIBHN_20035 | citrate synthase (unknown stereospecificity) | * | | IS256 | IS256 | | | | | |
| 4447835 | CIIBHN_20145 | ABC transporter substrate-binding protein | S279T | A | | | | | | T | |
| 4534006 | CIIBHN_20530 | type III secretion system export apparatus subunit SctV (bcrD, yscV, escV) | D598Y | G | T | T | T | T | T | T | T |
| 4584761 | | Intergenic | | C | T | | | | | | |
| 4621010 | CIIBHN_20925 | ilvD: dihydroxy-acid dehydratase | ins323S | ATGC | | | | | ATGCTGC | | |
| 4670782 | CIIBHN_21155 | GntR family transcriptional regulator | Synonymous | G | A | | | | | | |
| 4851242 | CIIBHN_21875 | Phage-base-V domain-containing protein | I611N | T | A | | | | | | |
| 4906519 | CIIBHN_22115 | type VI secretion system baseplate subunit TssK | Q39P | A | | C | | | | | |
| 4979756 | CIIBHN_22415 | transporter substrate-binding domain-containing protein | Synonymous | G | | | | | A | | |
| 4988809 | CIIBHN_22460 | LacI family transcriptional regulator | Synonymous | C | | | | A | | | |
| 5043168 | CIIBHN_22690 | Cytoplasmic protein | L62R | A | | C | | | | | |
| 5174950 | | Intergenic | | C | | | T | | | | |
| 5330854 | CIIBHN_23950 | ABC transporter permease | R166H | G | | | | | A | | |
| 5346567 | CIIBHN_24015 | uroporphyrinogen decarboxylase | Synonymous | C | | | | | | | T |
| 5405591 | CIIBHN_24305 | anthranilate synthase component I | V492D | T | | A | | | | | |
| 5483236 | CIIBHN_24665 | ABC transporter | P158L | G | A | | | | | | |
| 5641551 | CIIBHN_25340 | Branched chain amino acid ABC transporter substrate-binding protein | Synonymous | T | C | C | C | C | C | C | C |
| 5681646 | CIIBHN_25545 | Carnitine dehydratase | S98A | T | G | | | | | | |
| 5691723 | CIIBHN_25585 | Short-chain dehydrogenase | A69V | G | | | | | | | A |
| 5709159 | CIIBHN_25665 | Thymidylate kinase | E100K | G | A | A | A | A | A | A | A |
| 5709579 | CIIBHN_25665 | Thymidylate kinase | N240D | A | G | G | G | G | G | G | G |
| 5726613 | | Intergenic | | A | | C | | | | C | |
| 5744756 | CIIBHN_25850 | 50S ribosomal protein L24 | P49R | G | | | | | C | | |
| 5771137 | CIIBHN_25995 | DNA-directed RNA polymerase subunit beta' | P1346L | G | A | A | A | A | A | A | A |
| 5774839 | CIIBHN_25995 | DNA-directed RNA polymerase subunit beta' | A112V | G | A | | | | | | |

### Day 8

| Position | Gene Code | Gene | Consequence | Reference | 3 | 5 | 6 | 8 | 2 | 4 | 7 | 1 | Strain |
|---|---|---|---|---|---|---|---|---|---|---|---|---|---|
| | | | | | 2 | 2 | 2 | 2 | 2 | 2 | 2 | 2 | Cluster |
| | | | | | 1.8378 | 1.516 | 1.4324 | 1.2059 | 0.7568 | 0.5882 | 0.4865 | 0 | Median EOP |
| 10092 | | Intergenic | | A | | | | | | | | T | |
| 167924 | CIIBHN_00745 | Glutathione S-transferase | P81L | G | | | | | | A | | | |
| 206787 | CIIBHN_00950 | RNA polymerase sigma factor RpoH | Synonymous | C | | | G | | | | | | |
| 208884 | CIIBHN_00960 | Sodium:proton exchanger 208731..211241 | Frameshift AA 52/836 (84*) | GC | | GCC | GCC | | | | | | |
| 236439 | | Intergenic | | T | | | | C | C | | | | |
| 363559 | | Intergenic | | T | A | A | A | A | A | A | A | A | |
| 433860 | CIIBHN_01920 | Lipoprotein | Synonymous | T | | | | | | | | C | |
| 469553 | CIIBHN_02070 | DNA helicase | D29E | A | | | C | | | | | | |
| 542046 | CIIBHN_02435 | META domain-containing protein | Synonymous | T | C | C | C | C | C | C | C | C | |
| 579697 | | Intergenic | | T | C | C | C | | | | C | C | |
| 579703 | | Intergenic | | C | | | A | | | | | | |
| 609961 | | Intergenic | | G | | | | A | A | | | | |
| 658050 | CIIBHN_02920 | Malate dehydrogenase | Synonymous | C | | | | | | T | | | |
| 683477.. 683944 | CIIBHN_03020 | HTH tetR-type domain-containing protein (transcriptional regulator) | * | | IS481 | IS481 | IS481 | IS481 | | IS481 | IS481 | | |
| 684990.. 686186 | CIIBHN_03030 | LysR substrate-binding domain-containing protein (transcriptional regulator) | * | | IS481 | IS481 | IS481 | IS481 | | IS481 | IS481 | | |
| 729771 | CIIBHN_03240 | LysR family transcriptional regulator | Synonymous | C | | | | | | T | | | |
| 742208 | | Intergenic | | G | | | | | T | | | | |
| 907334 | CIIBHN_04070 | Carbon-phosphorus lyase complex subunit PhnI | S320I | G | | | | | T | | | | |
| 933320 | CIIBHN_04205 | Transcriptional regulator, possible shikimate kinase | Frameshift AA 27/301 (164*) | CCGC | | | CC | | | | | | |
| 933348 | CIIBHN_04205 | Transcriptional regulator, possible shikimate kinase | C36R | T | C | C | C | C | C | C | C | C | |
| 933419 | CIIBHN_04205 | Transcriptional regulator, possible shikimate kinase | Frameshift AA 59/301 (91*) | TG | | | | T | | | | | |
| 1092510 | | Intergenic | | T | A | A | A | | | A | A | A | |
| 1549571.. 1550377 | CIIBHN_06985 | RsxB: Electron transport complex subunit | * | | | | | Deleted | | | | | |
| 1578550 | CIIBHN_07120 | Phosphate acyltransferase | D215E | C | | A | | | | | | | |
| 1639769 | CIIBHN_07415 | Xaa-Pro aminopeptidase | Synonymous | C | G | | | | | | | | |
| 1703774 | CIIBHN_07715 | endopeptidase La | K392E | T | C | C | C | | | C | C | C | |
| 1823405 | CIIBHN_08215 | 2-aminoadipate aminotransferase, lysN | Q466R | T | C | C | C | C | C | C | C | C | |
| 1824452 | CIIBHN_08215 | 2-aminoadipate aminotransferase, lysN | D117V | T | | | | A | A | | | | |
| 1928888 | CIIBHN_08720 | LacI family transcriptional regulator | A75T | C | | | | | | G | | | |
| 2029057 | | Intergenic | | G | | | | | | | | A | |
| 2034824 | CIIBHN_09230 | MucB-RseB domain-containing protein | I153V | A | G | G | G | G | G | G | G | G | |
| 2217187 | CIIBHN_09965 | wecE: dTDP-4-amino-4,6-dideoxygalactose transaminase 2217081..2217962 | Frameshift AA 259/293 | TGG | | TG | | | | | | TG | |
| 2263781 | CIIBHN_10130 | MmgE/PrpD family protein | L250F | C | T | T | | | | | T | T | |
| 2331965.. 2333854 | CIIBHN_10415 | methyl-accepting chemotaxis protein I, serine sensor receptor (KEGG Ortholog of tsr) | * | | | Deleted | | | | | Deleted | | |
| 2382695 | CIIBHN_10670 | outer membrane protein assembly factor BamA | T518R | C | G | G | G | | | G | G | G | |
| 2392858 | CIIBHN_10720 | HTH araC/xylS-type domain-containing protein | V217G | A | | | | | C | | | | |
| 2439802 | CIIBHN_10965 | rcsC: regulator of capsular synthesis sensor histidine kinase | Q52* | C | T | T | T | T | T | T | T | T | |
| 2459392 | CIIBHN_11040 | TetR family transcriptional regulator | G239A | G | | | | | | | C | | |
| 2518442 | | Intergenic | | A | | | | C | | | | | |
| 2565287.. 2566870 | CIIBHN_11535 | EmrB/QacA family drug resistance transporter | * | | | | | | Deleted | | | | |
| 2630309 | CIIBHN_11785 | DUF802 domain-containing protein | Synonymous | G | C | C | | | | | C | C | |
| 2661550 | | Intergenic | | C | | | | T | | | | | |
| 3154590 | CIIBHN_14130 | putative chorismate pyruvate-lyase | Synonymous | C | | | | T | | | | | |
| 3376565 | CIIBHN_15100 | hypothetical protein | G185S | G | | | | A | A | | | | |
| 3419008 | CIIBHN_15280 | wzb: low molecular weight protein-tyrosine-phosphatase | T123A | T | | | | C | | | | | |
| 3419196 | CIIBHN_15280 | wzb: low molecular weight protein-tyrosine-phosphatase | A60V | G | A | A | | | | | | | |
| 3622057 | CIIBHN_16190 | Spermidine/putrescine ABC transporter permease | Synonymous | G | | | | | | | | A | |
| 3644402 | CIIBHN_16295 | alanine racemase | P73S | G | | | A | | | | | | |
| 3731250 | CIIBHN_16695 | 3-oxoacyl-ACP reductase | Synonymous | G | | | T | | | | | | |
| 3742189… 3787052 | CIIBHN_16750… CIIBHN_17130 | Prophage | * | | | | | | | Deleted | | | |
| 3853290 | CIIBHN_17470 | prephenate dehydratase | Frameshift (No*) AA 329/362 | CA | | | | | | | CAA | | |
| 3853293 | CIIBHN_17470 | prephenate dehydratase | Frameshift (No*) AA 329/362 | TA | | | | | | | | T | |
| 3854040 | CIIBHN_17470 | prephenate dehydratase | W80C | C | A | A | | | | | | | |
| 3854242 | CIIBHN_17470 | prephenate dehydratase | R13H | C | | | | | | T | | | |

### Day 8

| | | | | | | | | | | | | |
|---|---|---|---|---|---|---|---|---|---|---|---|---|
| 3857801 | CIIBHN_17480 | DNA gyrase subunit A | N87Y | T | | | | A | A | | | |
| 3937197 | CIIBHN_17970 | Chemotaxis protein cheW<br>(KEGG Ortholog of WspB) | A80V | C | | | | | | | T | |
| 3999174 | CIIBHN_18155 | citrate synthase | E410D | C | G | G | G | G | G | G | G | G |
| 4108696 | CIIBHN_18625 | Outer membrane porin protein BP0840, OmpP | T87S | T | A | A | A | A | A | A | A | A |
| 4223339 | CIIBHN_19195 | Type VI secretion protein | Synonymous | G | | | C | | | | | |
| 4328013 | | Intergenic | | A | G | G | G | G | G | G | G | G |
| 4328024 | | Intergenic | | G | A | A | A | A | A | A | A | A |
| 4358566 | CIIBHN_19750 | MFS transporter | A45D | C | | | | | | | A | |
| 4364236 | CIIBHN_19780 | hypothetical protein | M37V | A | G | G | G | G | G | G | G | G |
| 4368810 | CIIBHN_19795 | Amino acid deaminase | K357E | A | G | G | G | G | G | G | G | G |
| 4408497 | | Intergenic | | T | | | | | | A | | |
| 4420806..<br>4421057 | CIIBHN_20025 | hypothetical protein<br>(truncated filamentous haemagglutinin N-terminal domain) | * | | | | IS256 | | | | | |
| 4424326..<br>4425633 | CIIBHN_20035 | citrate synthase (unknown stereospecificity) | * | | | | IS256 | | | | | |
| 4524507 | CIIBHN_20475 | HrpE/YscL family type III secretion apparatus protein | Synonymous | G | | T | | | | | | |
| 4534006 | CIIBHN_20530 | type III secretion system export apparatus subunit SctV (bcrD, yscV, escV) | D598Y | G | T | T | T | T | T | T | T | T |
| 4584761 | | Intergenic | | C | | | | T | T | | | |
| 4670782 | CIIBHN_21155 | GntR family transcriptional regulator | Synonymous | G | | | | A | A | | | |
| 4697533 | CIIBHN_21255 | Hydrolase | A21V | C | | | | T | | | | |
| 4851242 | CIIBHN_21875 | Phage-base-V domain-containing protein | I611N | T | | | | A | A | | | |
| 5216167 | CIIBHN_21875 | Phage-base-V domain-containing protein | P178S | G | | | A | | | | | |
| 5349118 | CIIBHN_24025 | PepSY domain-containing protein | Synonymous | C | | | | | T | | | |
| 5375038 | CIIBHN_24155 | Amidohydro-rel domain-containing protein | C9S | A | | | | | | | T | |
| 5483236 | CIIBHN_24665 | ABC transporter | P158L | G | | | | A | A | | | |
| 5641551 | CIIBHN_25340 | Branched chain amino acid ABC transporter substrate-binding protein | Synonymous | T | C | C | C | C | C | C | C | C |
| 5676797 | CIIBHN_25515 | ABC transporter substrate-binding protein | R264Q | G | | | A | | | | | |
| 5681646 | CIIBHN_25545 | Carnitine dehydratase | S98A | T | | | | G | G | | | |
| 5709159 | CIIBHN_25665 | Thymidylate kinase | E100K | G | A | A | A | A | A | A | A | A |
| 5709579 | CIIBHN_25665 | Thymidylate kinase | N240D | A | G | G | G | G | G | G | G | G |
| 5726622 | | Intergenic | | G | | | | | | C | | |
| 5771137 | CIIBHN_25995 | DNA-directed RNA polymerase subunit beta' | P1346L | G | A | A | A | A | A | A | A | A |
| 5774839 | CIIBHN_25995 | DNA-directed RNA polymerase subunit beta' | A112V | G | | | | A | A | | | |

Day 17

| Position | Gene Code | Gene | Consequence | Reference | 1 | 2 | 3 | 4 | 5 | 6 | 7 | 8 | Strain |
|---|---|---|---|---|---|---|---|---|---|---|---|---|---|
| | | | | | 1 | 2 | 2 | 2 | 2 | 2 | 2 | 2 | Cluster |
| | | | | | 1.3188 | 0.8382 | 0.7246 | 0.493 | 0.286 | 0.2206 | 0 | 0 | Median EOP |
| 76633 | CIIBHN_00340 | Mandelate racemase | L218V | G | | | | | C | | | | |
| 151848..<br>153425 | CIIBHN_00675 | rhlE: ATP-dependent RNA helicase | * | | | | | | | | Deleted | | |
| 208884 | CIIBHN_00960 | Sodium:proton exchanger | Frameshift<br>AA 52/836 (84*) | GC | | | GCC | | | | | | |
| 228455..<br>229177 | CIIBHN_01060 | Rick-17kDa-Anti domain-containing protein | * | | Deleted | | | | | | Deleted | | |
| 236439 | | Intergenic | | T | | C | | | | | | | |
| 363559 | | Intergenic | | T | | A | A | A | A | A | A | A | |
| 542046 | CIIBHN_02435 | META domain-containing protein | Synonymous | T | | C | C | C | C | C | C | C | |
| 579697 | | Intergenic | | T | | | | C | C | C | | | |
| 609961 | | Intergenic | | G | | A | | | | | | | |
| 658050 | CIIBHN_02920 | Malate dehydrogenase | Synonymous | C | | | T | | | | T | | |
| 661248 | CIIBHN_02935 | Hydrolase | E107* | G | T | | | | | | | | |
| 683477..<br>683944 | CIIBHN_03020 | HTH tetR-type domain-containing protein<br>(transcriptional regulator) | * | | | | IS481 | IS481 | | IS481 | IS481 | IS481 | |
| 684990..<br>686186 | CIIBHN_03030 | LysR substrate-binding domain-containing protein<br>(transcriptional regulator) | * | | | | IS481 | IS481 | | IS481 | IS481 | IS481 | |
| 820797 | CIIBHN_03655 | histidine kinase | E222D | G | | T | | | | | | | |
| 933348 | CIIBHN_04205 | Transcriptional regulator, possible shikimate kinase | C36R | T | | C | C | C | C | C | C | C | |
| 933525 | CIIBHN_04205 | Transcriptional regulator, possible shikimate kinase | E95* | G | | | T | | | | T | | |
| 938293 | CIIBHN_04220 | benzoyl-CoA 2,3-epoxidase subunit BoxA | P191L | C | T | | | | | | | | |
| 968835 | CIIBHN_04355 | hypothetical protein | P737Q | G | | T | | | | | | | |
| 1092510 | | Intergenic | | T | | | A | A | A | A | A | A | |
| 1549571..<br>1550377 | CIIBHN_06985 | rsxB: Electron transport complex subunit | * | | | | | | | | Deleted | | |
| 1662441 | CIIBHN_07510 | General stress protein | Q61R | A | | | | | | G | | | |
| 1687807..<br>1688760 | CIIBHN_07640 | OmpR/PhoB-type domain-containing protein | * | | | | Deleted | | | | Deleted | | |
| 1703774 | CIIBHN_07715 | endopeptidase La | K392E | T | | | C | C | C | C | C | C | |
| 1823405 | CIIBHN_08215 | 2-aminoadipate aminotransferase, lysN | Q466R | T | | C | C | C | C | C | C | C | |
| 1824452 | CIIBHN_08215 | 2-aminoadipate aminotransferase, lysN | D117V | T | | A | | | | | | | |
| 2034824 | CIIBHN_09230 | MucB-RseB domain-containing protein | I153V | A | G | G | G | G | G | G | G | G | |
| 2042636 | CIIBHN_09265 | Carbon monoxide dehydrogenase subunit G | S140R | T | | | G | | | | | | |
| 2217187 | CIIBHN_09965 | wecE: dTDP-4-amino-4,6-dideoxygalactose transaminase<br>2217081..2217962 | Frameshift<br>AA 259/293 | TGG | | | | TG | | | | | |
| 2217485 | CIIBHN_09965 | wecE: dTDP-4-amino-4,6-dideoxygalactose transaminase | A160T | C | | | | | | T | | | |
| 2263781 | CIIBHN_10130 | MmgE/PrpD family protein | L250F | C | | | | T | T | T | | T | |
| 2349320..<br>2350732 | CIIBHN_10515 | fliK flagellar hook-length control protein | * | | | | | | | | Deleted | | |
| 2382695 | CIIBHN_10670 | outer membrane protein assembly factor BamA | T518R | C | | | G | G | G | G | G | G | |
| 2439802 | CIIBHN_10965 | rcsC: regulator of capsular synthesis sensor histidine kinase | Q52* | C | | T | T | T | T | T | T | T | |
| 2465756 | CIIBHN_11070 | 3-isopropylmalate dehydrogenase | E116* | G | | | | | | | T | | |
| 2630309 | CIIBHN_11785 | DUF802 domain-containing protein | Synonymous | G | | | | C | C | C | | C | |
| 2706760 | CIIBHN_12140 | Hydrolase | Synonymous | C | A | | | | | | | | |
| 3061299..<br>3062219 | CIIBHN_13725 | ANK-REP-REGION domain-containing protein | * | | | | | | | | Deleted | | |
| 3179158 | | Intergenic | | A | G | | | | | | | | |
| 3201814 | | Intergenic | | G | | | A | | | | | | |
| 3242255 | CIIBHN_14480 | Zinc-type alcohol dehydrogenase-like protein | Synonymous | G | | | | | | | | C | |
| 3376565 | CIIBHN_15100 | hypothetical protein | G185S | G | | A | | | | | | | |
| 3666766 | CIIBHN_16405 | Serine/threonine protein phosphatase | N124I | T | | A | | | | | | | |
| 3854040 | CIIBHN_17470 | prephenate dehydratase | W80C | C | | | | A | | A | | | |
| 3857801 | CIIBHN_17480 | DNA gyrase subunit A | N87Y | T | | A | | | | | | | |
| 3861740 | CIIBHN_17505 | Abasic site processing protein | F114L | C | | A | | | | | | | |
| 3926104 | CIIBHN_17915 | 3-deoxy-7-phosphoheptulonate synthase | M235I | G | | | | | | | | C | |
| 3937197 | CIIBHN_17970 | Chemotaxis protein cheW<br>(KEGG Ortholog of WspB) | A80V | C | | | | | T | | | | |
| 3999174 | CIIBHN_18155 | citrate synthase | E410D | C | | G | G | G | G | G | G | G | |
| 3990368..<br>3991603 | CIIBHN_18130 | odhB: 2-oxoglutarate dehydrogenase complex<br>dihydrolipoyllysine-residue succinyltransferase | * | | | | | | | | Deleted | | |
| 4042855 | CIIBHN_18340 | formate dehydrogenase-N subunit alpha | Synonymous | G | | | | | | | | A | |
| 4077437..<br>4078498 | CIIBHN_18490 | Type-4 uracil-DNA glycosylase | * | | | | | | | | Deleted | | |
| 4108696 | CIIBHN_18625 | Outer membrane porin protein BP0840, OmpP | T87S | T | | A | A | A | A | A | A | A | |
| 4121660 | CIIBHN_18670 | Betaine-aldehyde dehydrogenase | I143L | T | | | | | | | | G | |
| 4328013 | | Intergenic | | A | | G | G | G | G | G | G | G | |

Day 17

| | | | | | | | | | | | | |
|---|---|---|---|---|---|---|---|---|---|---|---|---|
| 4328024 | | Intergenic | | G | | A | A | A | A | A | A | A |
| 4364236 | CIIBHN_19780 | hypothetical protein | M37V | A | G | G | G | G | G | G | G | G |
| 4368810 | CIIBHN_19795 | Amino acid deaminase | K357E | A | | G | G | G | G | G | G | G |
| 4420806..<br>4421057 | CIIBHN_20025 | hypothetical protein<br>(truncated filamentous haemagglutinin N-terminal domain) | * | | | | | | | IS256 | | |
| 4424326..<br>4425633 | CIIBHN_20035 | citrate synthase (unknown stereospecificity) | * | | | | | | | IS256 | | |
| 4534006 | CIIBHN_20530 | type III secretion system export apparatus subunit SctV | D598Y | G | | T | T | T | T | T | T | T |
| 4577036 | CIIBHN_20740 | tripartite tricarboxylate transporter substrate binding protein<br>4576521..4577543 | Frameshift<br>AA 169/340 (169*) | TG | | | | | | | T | |
| 4584761 | | Intergenic | | C | | T | | | | | | |
| 4670782 | CIIBHN_21155 | GntR family transcriptional regulator | Synonymous | G | | A | | | | | | |
| 4680268 | CIIBHN_21190 | Methanethiol oxidase | G230D | C | | | | | T | | | |
| 4680906 | CIIBHN_21190 | Methanethiol oxidase | Synonymous | C | | | | | G | | | |
| 4708362 | CIIBHN_21295 | aconitate hydratase AcnA | L639Q | T | | | | | | | | A |
| 4851242 | CIIBHN_21875 | Phage-base-V domain-containing protein | I611N | T | | A | | | | | | |
| 4948771 | CIIBHN_22285 | Alpha/beta hydrolase | G123S | G | A | | | | | | | |
| 5284026 | CIIBHN_23760 | ATP-binding protein<br>5283991..5284848 | Frameshift (No*)<br>AA 17/285 | GCCCCCC | | | | GCCCCCCC | | | | |
| 5483236 | CIIBHN_24665 | ABC transporter | P158L | G | | A | | | | | | |
| 5641551 | CIIBHN_25340 | Branched chain amino acid ABC transporter substrate-binding protein | Synonymous | T | | C | C | C | C | C | C | C |
| 5681646 | CIIBHN_25545 | Carnitine dehydratase | S98A | T | | G | | | | | | |
| 5709159 | CIIBHN_25665 | Thymidylate kinase | E100K | G | | A | A | A | A | A | A | A |
| 5709579 | CIIBHN_25665 | Thymidylate kinase | N240D | A | | G | G | G | G | G | G | G |
| 5771137 | CIIBHN_25995 | DNA-directed RNA polymerase subunit beta' | P1346L | G | | A | A | A | A | A | A | A |
| 5774839 | CIIBHN_25995 | DNA-directed RNA polymerase subunit beta' | A112V | G | | A | | | | | | |

| Pos | Gene Number | Gene | Consequence | Reference | 1 | 2 | 3 | 4 | 5 | 6 | 7 | Strain |
|---|---|---|---|---|---|---|---|---|---|---|---|---|
| | | | | | 2 | 2 | 2 | 2 | 2 | 2 | 2 | Cluster |
| | | | | | 2.0541 | 2.0294 | 1.4595 | 1.371 | 1.2162 | 1.0541 | 0 | Median EOP |
| 228455..229177 | CIIBHN_01060 | Rick-17kDa-Anti domain-containing protein | * | | | | | | | Deleted | | |
| 236439 | | Intergenic | | T | | | | | | | C | |
| 363559 | | Intergenic | | T | A | A | A | A | A | A | A | |
| 393954 | | Intergenic | | T | | | | | | A | | |
| 542046 | CIIBHN_02435 | META domain-containing protein | Synonymous | T | C | C | C | C | C | C | C | |
| 579697 | | Intergenic | | T | C | C | | C | C | | | |
| 601884 | CIIBHN_02665 | lipoprotein insertase outer membrane protein LolB | S161Y | C | | | A | | | | | |
| 609961 | | Intergenic | | G | | | | | | | A | |
| 650542 | CIIBHN_02890 | TRAP transporter small permease | Synonymous | G | | | | | T | | | |
| 658050 | CIIBHN_02920 | Malate dehydrogenase | Synonymous | C | | | | | | T | | |
| 683477..683944 | CIIBHN_03020 | HTH tetR-type domain-containing protein (transcriptional regulator) | * | | | | IS481 | IS481 | | IS481 | IS481 | |
| 684990..686186 | CIIBHN_03030 | LysR substrate-binding domain-containing protein (transcriptional regulator) | * | | | | IS481 | IS481 | | IS481 | IS481 | |
| 746968 | CIIBHN_03310 | Short-chain dehydrogenase | Synonymous | C | | | T | | | | | |
| 843971 | CIIBHN_03750 | protein-glutamate O-methyltransferase | V331A | A | | | | | G | | | |
| 933348 | CIIBHN_04205 | Transcriptional regulator, possible shikimate kinase | C36R | T | C | C | C | C | C | C | C | |
| 933400 | CIIBHN_04205 | Transcriptional regulator, possible shikimate kinase | L53R | T | | | G | | | | | |
| 1010525 | CIIBHN_04515 | NAD-dependent dehydratase | Synonymous | C | | | | | T | | | |
| 1083919 | CIIBHN_04870 | Aminotransferase | C333G | A | | | C | | | | | |
| 1092510 | | Intergenic | | T | A | A | | A | A | A | | |
| 1478488 | CIIBHN_06690 | MFS transporter | P55L | C | | | | | | T | | |
| 1687807..1688760 | CIIBHN_07640 | OmpR/PhoB-type domain-containing protein | * | | | Deleted | | | | | | |
| 1702870 | CIIBHN_07715 | endopeptidase La | V693G | A | | | C | | | | | |
| 1703774 | CIIBHN_07715 | endopeptidase La | K392E | T | C | C | | C | C | C | | |
| 1823405 | CIIBHN_08215 | 2-aminoadipate aminotransferase, lysN | Q466R | T | C | C | C | C | C | C | C | |
| 1824452 | CIIBHN_08215 | 2-aminoadipate aminotransferase, lysN | D117V | T | | | | | | | A | |
| 2034824 | CIIBHN_09230 | MucB-RseB domain-containing protein | I153V | A | G | G | G | G | G | G | G | |
| 2087095 | | Intergenic | | G | | | A | | | | | |
| 2206607 | CIIBHN_09920 | ABC transporter ATP-binding protein | L139R | A | | | C | | | | | |
| 2217187 | CIIBHN_09965 | wecE: dTDP-4-amino-4,6-dideoxygalactose transaminase 2217081..2217962 | Frameshift AA 259/293 | TGG | | | | TG | | | | |
| 2217706 | CIIBHN_09965 | wecE: dTDP-4-amino-4,6-dideoxygalactose transaminase | T86I | G | | | A | | | | | |
| 2218801 | CIIBHN_09970 | Putative S-adenosylmethionine-dependent methyltransferase | P25A | G | | | C | | | | | |
| 2263781 | CIIBHN_10130 | MmgE/PrpD family protein | L250F | C | T | T | | T | T | | | |
| 2277944 | CIIBHN_10185 | dihydrolipoyl dehydrogenase | D295N | G | | | | | | | A | |
| 2331965..2333854 | CIIBHN_10415 | methyl-accepting chemotaxis protein I, serine sensor receptor (KEGG Ortholog of tsr) | * | | Deleted | | Deleted | | | | | |
| 2333862..2334347 | CIIBHN_10420 | Outer membrane protein assembly factor BamE | * | | | | Deleted | | | | | |
| 2334439..2336184 | CIIBHN_10425 | methyl-accepting chemotaxis protein I, serine sensor receptor (KEGG Ortholog of tsr) | * | | | | Deleted | | | | | |
| 2382695 | CIIBHN_10670 | outer membrane protein assembly factor BamA | T518R | C | G | G | | G | G | G | | |
| 2439802 | CIIBHN_10965 | rcsC: regulator of capsular synthesis sensor histidine kinase | Q52* | C | T | T | T | T | T | T | T | |
| 2630309 | CIIBHN_11785 | DUF802 domain-containing protein | Synonymous | G | C | C | | C | C | | | |
| 2888609 | CIIBHN_12985 | acetylornithine deacetylase | S106N | G | | | | | | A | | |
| 2926396 | CIIBHN_13155 | 2-keto-4-pentenoate hydratase | A241V | C | | | | | | T | | |
| 3143021 | CIIBHN_14080 | 4-chlorobenzoate--CoA ligase | L55R | A | | | | | C | | | |
| 3376565 | CIIBHN_15100 | hypothetical protein | G185S | G | | | | | | | A | |
| 3419153 | CIIBHN_15280 | wzb: low molecular weight protein-tyrosine-phosphatase | Q74H | C | | | A | | | | | |
| 3419374 | CIIBHN_15280 | wzb: low molecular weight protein-tyrosine-phosphatase | Start-Loss M1L | T | | | | | A | | | |
| 3419409 | | Intergenic | | A | G | G | | G | | | | |
| 3557321 | CIIBHN_15910 | aminopeptidase N | T153M | C | | | | | | T | | |
| 3853290 | CIIBHN_17470 | prephenate dehydratase | Frameshift AA 330/362 | CA | | CAA | | | | | | |
| 3854116 | CIIBHN_17470 | prephenate dehydratase | R55H | C | | | | | T | | | |
| 3857801 | CIIBHN_17480 | DNA gyrase subunit A | N87Y | T | | | | | | | A | |
| 3937197 | CIIBHN_17970 | Chemotaxis protein cheW (KEGG Ortholog of WspB) | A80V | C | T | T | | T | | | | |
| 3999174 | CIIBHN_18155 | citrate synthase | E410D | C | G | G | G | G | G | G | G | |
| 4010101 | CIIBHN_18210 | Glutathione S-transferase | R180C | C | | T | | | | | | |
| 4108696 | CIIBHN_18625 | Outer membrane porin protein BP0840, OmpP | T87S | T | A | A | A | A | A | A | A | |
| 4328013 | | Intergenic | | A | G | G | G | G | G | G | G | |

| | | | | | | | | | | | |
|---|---|---|---|---|---|---|---|---|---|---|---|
| 4328024 | Intergenic | | | G | A | A | A | A | A | A | A |
| 4364236 | CIIBHN_19780 | hypothetical protein | M37V | A | G | G | G | G | G | G | G |
| 4368810 | CIIBHN_19795 | Amino acid deaminase | K357E | A | G | G | G | G | G | G | G |
| 4420806..<br>4421057 | CIIBHN_20025 | hypothetical protein<br>(truncated filamentous haemagglutinin N-terminal domain) | * | | | IS256 | IS256 | | | | |
| 4424326..<br>4425633 | CIIBHN_20035 | citrate synthase (unknown stereospecificity) | * | | | IS256 | IS256 | | | | |
| 4534006 | CIIBHN_20530 | type III secretion system export apparatus subunit SctV | D598Y | G | T | T | T | T | T | T | T |
| 4584761 | Intergenic | | | C | | | | | | | T |
| 4670782 | CIIBHN_21155 | GntR family transcriptional regulator | Synonymous | G | | | | | | | A |
| 4851242 | CIIBHN_21875 | Phage-base-V domain-containing protein | I611N | T | | | | | | | A |
| 4875080 | CIIBHN_21960 | ROK family protein | T333N | C | A | A | | A | | | |
| 5253635 | CIIBHN_23600 | Outer membrane lipoprotein Blc | A23S | G | | | | | T | | |
| 5280560 | Intergenic | | | G | | | | A | | | |
| 5483236 | CIIBHN_24665 | ABC transporter | P158L | G | | | | | | | A |
| 5641551 | CIIBHN_25340 | Branched chain amino acid ABC transporter substrate-binding protein | Synonymous | T | C | C | C | C | C | C | C |
| 5681646 | CIIBHN_25545 | Carnitine dehydratase | S98A | T | | | | | | | G |
| 5709159 | CIIBHN_25665 | Thymidylate kinase | E100K | G | A | A | A | A | A | A | A |
| 5709579 | CIIBHN_25665 | Thymidylate kinase | N240D | A | G | G | G | G | G | G | G |
| 5771137 | CIIBHN_25995 | DNA-directed RNA polymerase subunit beta' | P1346L | G | A | A | A | A | A | A | A |
| 5774839 | CIIBHN_25995 | DNA-directed RNA polymerase subunit beta' | A112V | G | | | | | | | A |

| Pos | Gene Number | Gene | Consequence | Reference | 1 | 2 | 3 | 4 | 5 | 6 | Strain |
|---|---|---|---|---|---|---|---|---|---|---|---|
| | | | | | 1 | 1 | 1 | 1 | 2 | 1 | Cluster |
| | | | | | 1.7183 | 1.3661 | 1.2676 | 1.1831 | 1.0423 | 0.837 | Median EOP |
| 208884 | CIIBHN_00960 | Sodium:proton exchanger | Frameshift AA 52/836 (84*) | GC | | | | | GCC | | |
| 251903 | CIIBHN_01175 | 3-dmu-9-3-mt domain-containing protein | P66R | C | | G | | | | | |
| 363559 | | Intergenic | | T | | | | | A | | |
| 542046 | CIIBHN_02435 | META domain-containing protein | Synonymous | T | | | | | C | | |
| 658050 | CIIBHN_02920 | Malate dehydrogenase | Synonymous | C | | | | | T | | |
| 683477.. 683944 | CIIBHN_03020 | HTH tetR-type domain-containing protein (transcriptional regulator) | * | | IS481 | IS481 | | IS481 | IS481 | | |
| 684990.. 686186 | CIIBHN_03030 | LysR substrate-binding domain-containing protein (transcriptional regulator) | * | | IS481 | IS481 | | IS481 | IS481 | | |
| 742212 | | Intergenic | | G | | A | | | | | |
| 933348 | CIIBHN_04205 | Transcriptional regulator, possible shikimate kinase | C36R | T | | | | | C | | |
| 1092510 | | Intergenic | | T | | | | | A | | |
| 1383568 | CIIBHN_06260 | 4-hydroxy-tetrahydrodipicolinate synthase | S112F | C | | | T | | | | |
| 1484575 | CIIBHN_06725 | histidine kinase | Synonymous | G | | | | | T | | |
| 1549571.. 1550377 | CIIBHN_06985 | rsxB: Electron transport complex subunit | * | | | | | | Deleted | | |
| 1588330 | CIIBHN_07175 | translation elongation factor 4 | Synonymous | C | A | | | | | A | |
| 1703774 | CIIBHN_07715 | endopeptidase La | K392E | T | | | | | C | | |
| 1823405 | CIIBHN_08215 | 2-aminoadipate aminotransferase, lysN | Q466R | T | | | | | C | | |
| 1870339 | | Intergenic | | C | | | | | T | | |
| 2034824 | CIIBHN_09230 | MucB-RseB domain-containing protein | I153V | A | G | G | G | G | G | G | |
| 2331965.. 2333854 | CIIBHN_10415 | methyl-accepting chemotaxis protein I, serine sensor receptor (KEGG Ortholog of tsr) | * | | | | Deleted | | Deleted | | |
| 2382695 | CIIBHN_10670 | outer membrane protein assembly factor BamA | T518R | C | | | | | G | | |
| 2439802 | CIIBHN_10965 | rcsC: regulator of capsular synthesis sensor histidine kinase | Q52* | C | | | | | T | | |
| 2505447 | CIIBHN_11255 | Short-chain dehydrogenase | Synonymous | G | | | | | | A | |
| 2518442 | | Intergenic | | A | | | | | C | | |
| 2538712 | CIIBHN_11415 | CAAX protease | G166S | G | | | | A | | | |
| 2695873 | CIIBHN_12085 | MFS transporter | A235P | C | | | | | | G | |
| 2882051.. 2882707 | CIIBHN_12950 | hypothetical protein | * | | | IS256 | | | | | |
| 3036263 | CIIBHN_13620 | YicC family protein | Synonymous | C | | T | | | | | |
| 3352807 | CIIBHN_14995 | NmrA family transcriptional regulator | A71P | C | | | | | | G | |
| 3418943 | CIIBHN_15280 | wzb: low molecular weight protein-tyrosine-phosphatase | D144E | G | | | | | C | | |
| 3421865 | CIIBHN_15290 | Aminotransferase | R28W | C | | | | | T | | |
| 3773048 | | Intergenic | | G | | | T | | | | |
| 3819881 | CIIBHN_17305 | phenylacetic acid degradation operon negative regulatory protein PaaX | R86W | C | | | | | T | | |
| 3925517 | CIIBHN_17915 | 3-deoxy-7-phosphoheptulonate synthase | A40T | G | | | | A | | | |
| 3999174 | CIIBHN_18155 | citrate synthase | E410D | C | | | | | G | | |
| 4040180.. 4041070 | CIIBHN_18335 | LysR family transcriptional regulator | * | | | | | IS481 | | | |
| 4041260.. 4044328 | CIIBHN_18340 | formate dehydrogenase-N subunit alpha | * | | | | | IS481 | | | |
| 4108696 | CIIBHN_18625 | Outer membrane porin protein BP0840, OmpP | T87S | T | | | | | A | | |
| 4188041 | CIIBHN_19020 | Tripartite tricarboxylate transporter substrate binding protein | Synonymous | C | T | | | | | T | |
| 4253880 | CIIBHN_19325 | Tyrosine protein kinase | Synonymous | G | | | | | A | | |
| 4270472 | CIIBHN_19390 | UDP-N-acetylglucosamine 2-epimerase | Synonymous | G | A | | | | | | |
| 4328013 | | Intergenic | | A | | | | | G | | |
| 4328024 | | Intergenic | | G | | | | | A | | |
| 4364236 | CIIBHN_19780 | hypothetical protein | M37V | A | G | G | G | G | G | G | |
| 4368810 | CIIBHN_19795 | Amino acid deaminase | K357E | A | | | | | G | | |
| 4420806.. 4421057 | CIIBHN_20025 | hypothetical protein (truncated filamentous haemagglutinin N-terminal domain) | * | | | | | IS256 | | | |
| 4424326.. 4425633 | CIIBHN_20035 | citrate synthase (unknown stereospecificity) | * | | | | | IS256 | | | |
| 4431709 | CIIBHN_20065 | Bcr/CflA family efflux transporter | R74Q | C | | | T | | | | |
| 4534006 | CIIBHN_20530 | type III secretion system export apparatus subunit SctV | D598Y | G | | | | | T | | |
| 4705834 | CIIBHN_21290 | Transcriptional regulator | Y119S | A | | C | | | | | |
| 5090622 | CIIBHN_22910 | anmK: anhydro-N-acetylmuramic acid kinase | Frameshift AA 43/372 | GCCC | GCCCC | | | | | GCCCC | |
| 5091449 | CIIBHN_22910 | anmK: anhydro-N-acetylmuramic acid kinase | L312R | T | | G | | | | | |
| 5139065 | CIIBHN_23125 | LPS export ABC transporter permease LptG | K168T | A | | C | | | | | |
| 5410801 | | Intergenic | | C | | | | G | | | |
| 5641551 | CIIBHN_25340 | Branched chain amino acid ABC transporter substrate-binding protein | Synonymous | T | | | | | C | | |
| 5709159 | CIIBHN_25665 | Thymidylate kinase | E100K | G | | | | | A | | |

| | | | | | | | | | |
|---|---|---|---|---|---|---|---|---|---|
| 5709579 | CIIBHN_25665 | Thymidylate kinase | N240D | A | | | | | G |
| 5771137 | CIIBHN_25995 | DNA-directed RNA polymerase subunit beta' | P1346L | G | | | | | A |
